# Lactoferricins access the cytosol of *Escherichia coli* within few seconds

**DOI:** 10.1101/2021.09.24.461681

**Authors:** Enrico F. Semeraro, Lisa Marx, Johannes Mandl, Ilse Letofsky-Papst, Claudia Mayrhofer, Moritz P. K. Frewein, Haden L. Scott, Sylvain Prévost, Helmut Bergler, Karl Lohner, Georg Pabst

## Abstract

We report the real-time response of *E. coli* to lactoferricin-derived antimicrobial peptides (AMPs) on length-scales bridging microscopic cell-sizes to nanoscopic lipid packing using millisecond time-resolved synchrotron small-angle X-ray scattering. Coupling a multi-scale scattering data analysis to biophysical assays for peptide partitioning revealed that the AMPs rapidly saturate the bacterial envelope and reach the cytosol within less than three seconds—much faster than previously considered. Final cytosolic AMP concentrations of ~ 100 mM suggest an efficient shut-down of metabolism as primary cause for bacterial killing. On the other hand, the damage of the cell envelope is a collateral effect of AMP activity that does not kill the bacteria. This implies that the impairment of the membrane barrier is a necessary but not sufficient condition for microbial killing by lactoferricins. The most efficient AMP studied exceeds others in both speed of reaching cytoplasm and lowest cytosolic peptide concentration.

## Introduction

Progress in designing antibiotics with novel key-lock mechanisms is not keeping pace with the worldwide growing number of (multi) resistant bacterial strains, encouraging significant research efforts in promising alternatives such as antimicrobial peptides (AMPs) (***Lohner, 2001***). AMPs are part of the natural innate immune system and provide a first line of defence against pathogens. Their advantage as compared to conventional antibiotics relies on a rapid impairment of the barrier-function of the bacterial envelope by unspecific physical interactions, often coupled to an ensuing targeting of bacterial DNA or ribosomes [for review see, e.g., ***Wimley and Hristova*** (***2011***); ***Lohner*** (***2017***); ***Malanovic et al.*** (***2020***)].

Membrane-active AMPs contain specific sequences of cationic and apolar amino-acids, granting high affinity to the hydrophobic core of lipid membranes and selectivity towards the negatively charged surfaces of bacterial envelopes. However, despite intense research for several decades, a comprehensive understanding of the specific series of events that pertain to the bactericidal or bacteriostatic activity of AMPs is still elusive. To large extent this is due to the persisting challenge of merging results from *in vitro* studies with those obtained from lipid membrane mimics, often leading to significant controversies (***Wimley and Hristova, 2011***). This is nurtured, on the one hand, by difficulties in engineering lipid model systems of sufficiently high complexity to mimic the diverse physicochemical properties of bacterial membranes. On the other hand, the complexity of live bacteria challenges experimental and computational techniques to obtain quantitative results on the molecular level. For example, cryogenic transmission electron microscopy (TEM) provides high sub-cellular spatial resolution, but might give misleading information due to artefacts that potentially originate from staining or invasive sample preparation. Moreover, structural kinetics occurring in the seconds time scale are yet not accessible to cryo-TEM on cells, but would be needed to unravel the sequence of events induced by AMP activity. High-speed atomic force microscopy, for example, showed that a corrugation of the outer surface of live bacteria occurred about as fast as the intrinsic time resolution of the experiment, i.e., within the first 13 seconds after addition of the AMP (***Fantner et al., 2010***). However, such experiments do not provide insight on concurring intracellular changes. Video fluorescence microscopy provides the appropriate spatiotemporal resolution to differentiate AMP activity in different cells within several tens of seconds [see ***Choi et al.*** (***2016***) for review]. By combining fluorescence labeling schemes for peptides or cellular content, it has been reported that peptides preferentially attack septating cells and often reach the cytoplasm within few minutes, suggesting that the final target for arresting bacterial growth or killing is not the cytoplasmic membrane (***Sochacki et al., 2011***). However, fluorescence labeling may easily tweak the delicate balance of macromolecular interactions and thus affect experimental observations.

We have recently reported an analytical model for elastic X-ray and neutron scattering from live *Escherichia coli* without the need to resort to any specific labelling technique (***Semeraro et al., 2021***). In particular, we combined the different sensitivities of X-rays and neutrons to matter, including H/D contrast variation, with a compositional multi-scale model. This allowed us to detail the bacterial hierarchical structure on lengths scales bridging four orders of magnitude, i.e., spanning from bacterial size to the molecular packing of lipopolysacherides (LPS) in the outer leaflet of the outer membrane. Here we use this model, taking advantage of the fact that the full breath of structural information is encoded in a single scattering pattern, and exploit millisecond time-resolved synchrotron (ultra) small-angle X-ray scattering (USAXS/SAXS) to study the response of *E. coli* to three lactoferricin-derived AMPs: LF11-215 (FWRIRIRR-NH_2_), LF11-324 (PFFWRIRIRR-NH_2_) and O-LF11-215 (octanoyl-FWRIRIRR-NH_2_). The activity of these AMPs has been studied before, both *in vitro* and in bacterial membrane mimics, using an array of biophysical and biochemical assays (***Zweytick et al., 2011, 2014***; ***Sánchez-Gómez et al., 2015***; ***Marx et al., 2021b***).

Joining these elastic scattering experiments with cryo-TEM and assays for determining peptide partitioning as a function of peptide activity enabled us to gain unprecedented insight into the peptide-induced sequence of events. Strikingly, we found that the studied peptides are able to reach the bacterial cytosol just within few seconds, much faster than previously reported (***Choi et al., 2016***). Concomitantly this leads to a jump of peptide concentration in the cytosol, reaching about 100 mM at full growth inhibition. The most effective AMP presently studied, LF11-324, excels others by an increased speed of translocation and lowest cytosolic concentration. We also observed collateral damage of the bacterial cell envelope (loss of LPS packing, loss of positional correlations between outer and inner membranes, vesiculation/tubulation, cell shrinkage) in agreement with previous studies (***Zweytick et al., 2011***). However, these changes occurred at later time points and also for peptide concentrations far below the minimum inhibitory concentration (MIC). The primary cause for bactericidal or bacteriostatic activity of the presently studied peptides is thus not a damage of the structural integrity of the cell-wall, but appears to be a fast and efficient shut-down of bacterial metabolic activity.

## Results

### Defining structural reference states of AMP activity in *E. coli*

Unravelling the time-line of structural events occurring in *E. coli* due to lactoferricin activity by USAXS/SAXS necessitates a detailed prior characterization of two reference states: (*i*) neat bacteria before AMP administration (‘initial-state’), and (*ii*) AMP-affected/killed bacteria (‘end-state’). Here, end-state refers to one hour of incubation of bacteria at a given AMP concentration. We have recently reported initial-state structures of different *E. coli* strains at different hierarchical length scales—including size of bacteria, distance between inner and outer membranes, and LPS packing density—in terms of a multi-scale analytical model using joint USAXS/SAXS and (very) small-angle neutron scattering (VSANS)/SANS experiments (***Semeraro et al., 2021***). The same concept was applied here to reveal the end-state structure of *E. coli*. In order to remove ambiguities in adjustable parameters due to ensemble averaging, these experiments where coupled to cryo-TEM (**Figure 1**). It is important to note that SAXS/SANS experiments require high bacterial concentrations (~ 10^9^ CFU/ml). Since this affects AMP activity (***Marx et al., 2021b***), we report the MIC as a function of cell number density; the corresponding MIC values for the presently studied AMPs are reported in **Figure 1–Figure Supplement 1**.

**Figure 1.**
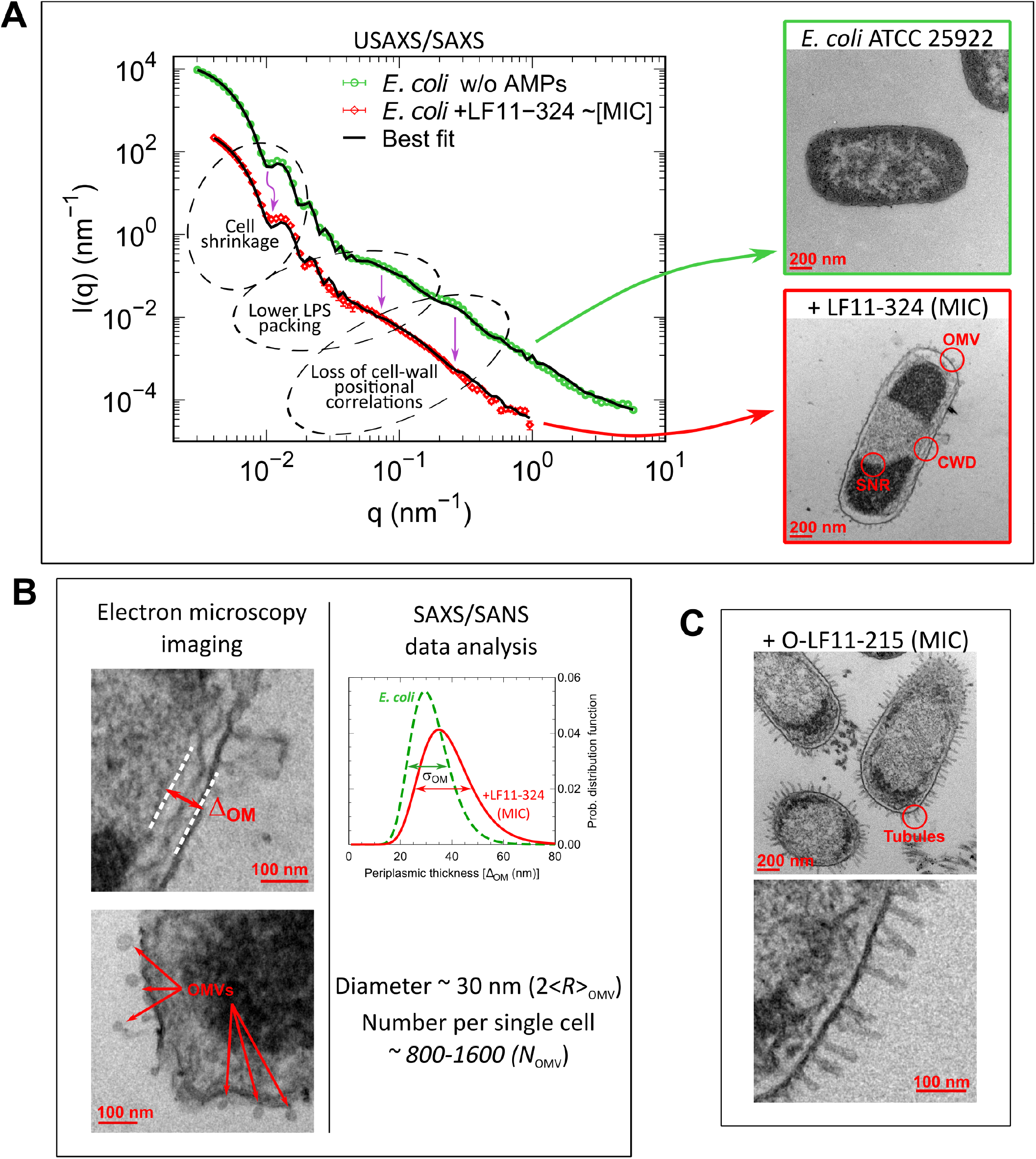
(**A**) Mapping the main structural changes in *E. coli* ATCC 25922 (green symbols) upon 1 h incubation with LF11-324 (red symbols) as observed by USAXS/SAXS and TEM. Scattering data of *E. coli* ATCC 25922 are from ***Semeraro et al.*** (***2021***). Black lines are the best fits using **Equation 6**. <u>Abbreviations:</u> OMV: outer membrane vesicle formation; CWD: cell-wall damaging; SNR: phase separation of the nucleoid region. (**B**) TEM examples of membrane detachment and OMV formation due to LF11-324, and respective *ensemble* results from scattering data analysis for the distance distribution between inner and outer membranes. (**C**) Bacteria upon 1 h incubation with O-LF11-215, showing the formation of tube-like protrusions. **Figure 1–Figure supplement 1.** Cell number-dependent MIC plots for different peptides. **Figure 1–Figure supplement 2.** Comparison between USAXS/SAXS and contrast-variation SANS data and details of the scattering data analysis. **Figure 1–Figure supplement 3.** TEM observations for LF11-215, LF11-324 and O-LF11-215 at the MICs and sub-MICs.

Small-angle scattering (SAS) patterns of initial and end-states showed distinct differences, many of which can be compared to cryo-TEM results. Membrane ruffling, for example, originating mainly from increased fluctuations of cytoplasmic membranes, leads to a modification of the inner/outer membrane distance distribution function (**Figure 1**A-B and **Figure 1–Figure Supplement 2**F). Shrinking of bacterial size, in turn is observed by changes in intensity modulation at very low scattering vectors. In addition, hidden to TEM, but revealed by USAXS/SAXS/SANS are changes to the lateral LPS density when focusing on the scattering shoulder at *q* ~ 0.07 nm^−1^. The lowering of its intensity might originate either from a lower LPS surface density (LPS packing), or a loss of positional correlations along the surface (membrane roughness or waving), or a combination both. The end-state scattering patterns of bacteria in the presence of the well distinct MICs of LF11-324 and LF11-215 were identical. The scattering patterns of O-LF11-215 instead indicated the formation of peptide aggregates (Appendix 1), which results from its increased hydrophobicity and hence lower critical aggregate concentration in buffer. LF11-324 and LF11-215 caused the formation of extracellular vesicles, also known as outer membrane vesicles (OMVs), clearly observed by TEM (**Figure 1**A-B) and with an average diameter of ~ 30 nm diameter as revealed by SAS data analyis. O-LF11-215, in contrast, causes the formation of extramembranous tubes (**Figure 1**C), to which USAXS/SAXS/SANS is not sensitive to. Further, the increased scattering contributions originating from O-LF11-215 aggregates impeded the detection of OMVs. Notably, SAS is insensitive to the inner cytosolic structure of bacteria (***Semeraro et al., 2021***). Hence the peptide induced separation of the nucleoid region from the nucleoid-free cytosol (**Figure 1**A) reported also previously from TEM (***Zweytick et al., 2011***), is not observed in our scattering data.

**Table 1** summarizes the changes between initial and end-states for LF11-324from the joint USAXS/SAXS and SANS multi-scale analysis. In particular, we report (*i*) contrast changes in terms of the scattering length densities (SLDs) of the cytoplasm, *ρ_CP_*, periplasm, *ρ_PP_*, and peptidoglycan, *ρ_PG_*, (*ii*) microscopic to mesoscopic structural size changes of bacteria, approximated by an ellipsoid of inner radius *R*, and distance between inner and outer membranes, Δ_OM_, as well as the corresponding distribution of distances between the two membranes, *σ*_OM_, and (*iii*) nanoscopic structural changes as observed for the average number of LPS, *N_OS_*, per cell. See ***Semeraro et al.*** (***2021***), for a justification of all used parameters. The decrease of *ρ*_CP_, along with the increase of *ρ*_PP_, signifies leakage of mainly low-weight molecules from the cytoplasm (**Figure 1–Figure Supplement 2**B-C). Notably, the observed cell shrinkage of ~ 5% leads to a decrease of cell surface of approximately 2 × 10^6^ nm^2^. Apparently this is at least in part compensated by OMV formation, as suggested by their total surface estimate of ~ (2 – 6) × 10^6^ nm^2^ from SAS analysis (see Appendix 2).

**Table 1.** Change of *E. coli* structure due to LF11-324 ([*P*] ~ MIC) as observed from USAXS/SAXS/SANS data analysis.

| Parameters | Values |
| --- | --- |
| $\Delta\rho_{CP} \times 10^{-4} \text{ (nm}^{-2}\text{)}$ | $-0.17 \pm 0.02^{(X)}$; $-0.14 \pm 0.05^{(N)}$ |
| $\Delta\rho_{PP} \times 10^{-4} \text{ (nm}^{-2}\text{)}$ | $0.18 \pm 0.06^{(X)}$; $0.12 \pm 0.04^{(N)}$ |
| $\Delta\Delta_{OM} \text{ (nm)}$ | $6 \pm 3$ |
| $\Delta\sigma_{OM} \text{ (nm)}$ | $3.4 \pm 1.7$ |
| $\Delta\rho_{PG} \times 10^{-4} \text{ (nm}^{-2}\text{)}$ | $-0.27 \pm 0.07^{(X)}$; $-0.59 \pm 0.14^{(N)}$ |
| $\Delta N_{OS} \times 10^6$ | $-2.0 \pm 0.4^{(X)}$; $-1.6 \pm 0.8^{(N)}$ |
| $\Delta R \text{ (nm)}$ | $-27 \pm 7$ |

(X) from SAXS. (N) from SANS (SLDs were obtained by extrapolating to 0 wt% D<sub>2</sub>O).

### Kinetics: time-resolved USAXS/SAXS

The structural transitions from initial to end-state were followed by USAXS/SAXS at millisecond time resolution. Stopped-flow mixing ensured thorough and rapid re-dispersion (mixing time of 50 ms) of peptides and bacteria (**Figure 2–Figure Supplement 1**) and led to immediate changes of scattering patterns. Firstly, LPS packing 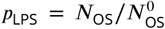 (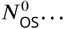 initial number of LPS per cell) started to decrease at Δ*t* ~ 10 s after mixing, independent of LF11-324 concentration, i.e. even at [*P*] = 0.3×MIC (**Figure 2**A). The loss of cytoplasmic content, as observed in *ρ*_CP_, commenced at similar times, although the increase of *ρ*_PP_ can be tracked down to 3 s for the highest LF11-324 concentration (**Figure 2**B). Also the drop of *R* exhibited concentration-dependent kinetics (**Figure 2**C), starting at 20–50 s for [*P*] = 1.2×MIC and 0.7×MIC, and > 10 min for [*P*] = 0.3×MIC. Changes of Δ_OM_, *σ*_OM_ and *ρ*_PG_ in turn seem to be largely decoupled from these early events, with an onset of 2–10 minutes after peptide addition (**Figure 2**D – F). Interestingly, LF11-215 led to very similar kinetics for all parameters, except for slower changes of *ρ*_CP_ and *ρ*_PP_ (**Figure 2–Figure Supplement 2**, **Figure 2–Figure Supplement 3**). This suggests that the permeability of the cell-wall is affected in a concentration-dependent and peptide-specific manner. The similar onsets for changes of Δ_OM_, *σ*_OM_ and *ρ*_PG_ in turn suggest that this does not apply to the overall stability of the cell envelope. Interestingly, no decrease of *p*_LPS_ was observed for LF11-215 at [*P*] = 1.6×MIC (**Figure 2–Figure Supplement 3**A). In turn, changes of *ρ*_CP_ and *ρ*_PP_ occurred for O-LF11-215 at about similar times than for LF11-324 (**Figure 2**B). Later onsets of changes were observed for *p*_LPS_ (~ 2 min, **Figure 2**G) and *R* (~10 min, **Figure 2**I) in this case. The initial increase of *p*_LPS_ is likely to be an artefact, probably due to the overlap of *I*_cell_(*q*) and *I*_clu_(*q*) (see **Figure 2–Figure Supplement 1**D). The analysis of end-states was obscured for O-LF11-215 due to rapid sample sedimentation triggered by macroscopic peptide/bacteria clustering.

**Figure 2.**
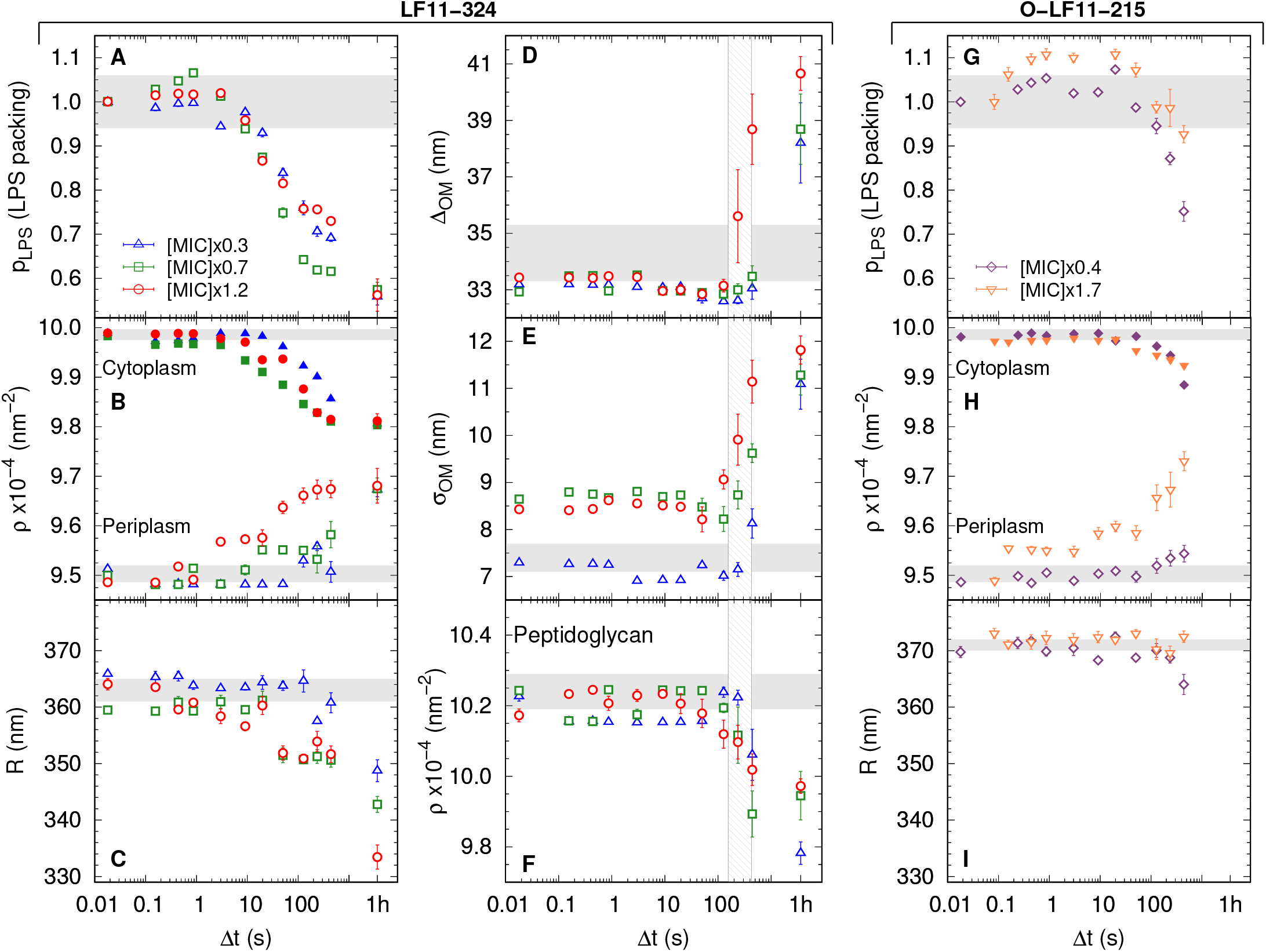
(**A-F**) Kinetics of the bacterial structural response to attack by LF11-324; results for three different peptide concentrations are shown. LPS packing (**A**); cytoplasm and periplasm SLDs (**B**); minor radius of the cell (**C**); intermembrane distance (~ periplasm thickness) (**D**) and its deviation (**E**); and peptidoglycan SLD (**F**). (**G-I**) Bacterial response to O-LF11-215 at two concentrations. LPS packing (**G**); cytoplasm and periplasm SLDs (**H**); and minor ellipsoidal radius of the cell (**I**). Thick gray bands mark the degree of confidence from bacterial systems w/o peptides [see **Table 1** and ***Semeraro et al.*** (***2021***)], except for (**C**) and (**I**), where they refer to the average of the current cell radii at △*t* = 0.0175 s. Fluctuations of initial values can be due to biological diversity. The vertical gray grid in (**D-F**) indicates the time range of cell-wall damage. Note that this range does not dependent on peptide concentration. Results at Δ*t* =1 hour refer to end-states, when available. **Figure 2–Figure supplement 1.** Schematic of the stopped-flow rapid mixing USAXS/SAXS experiments, including selected scattering patterns. **Figure 2–Figure supplement 2.** Kinetics of the adjustable parameters for LF11-215 and O-LF11-215 systems. **Figure 2–Figure supplement 3.** Kinetics of the adjustable parameters for LF11-215 systems.

Scattering originating from LF11-215/LF11-324-induced OMV formation was discernible for Δ*t* > 1 – 2 min. Yet, the average number of formed OMVs appear to be peptide- and concentration-independent within the first 10 min. Finally, the peptide cluster term, introduced for the analysis of O-LF11-215, enabled us to estimate that a large increase of peptide uptake starts after about 2 min. No dependence on peptide concentration was observed (**Figure 3**B).

**Figure 3.**
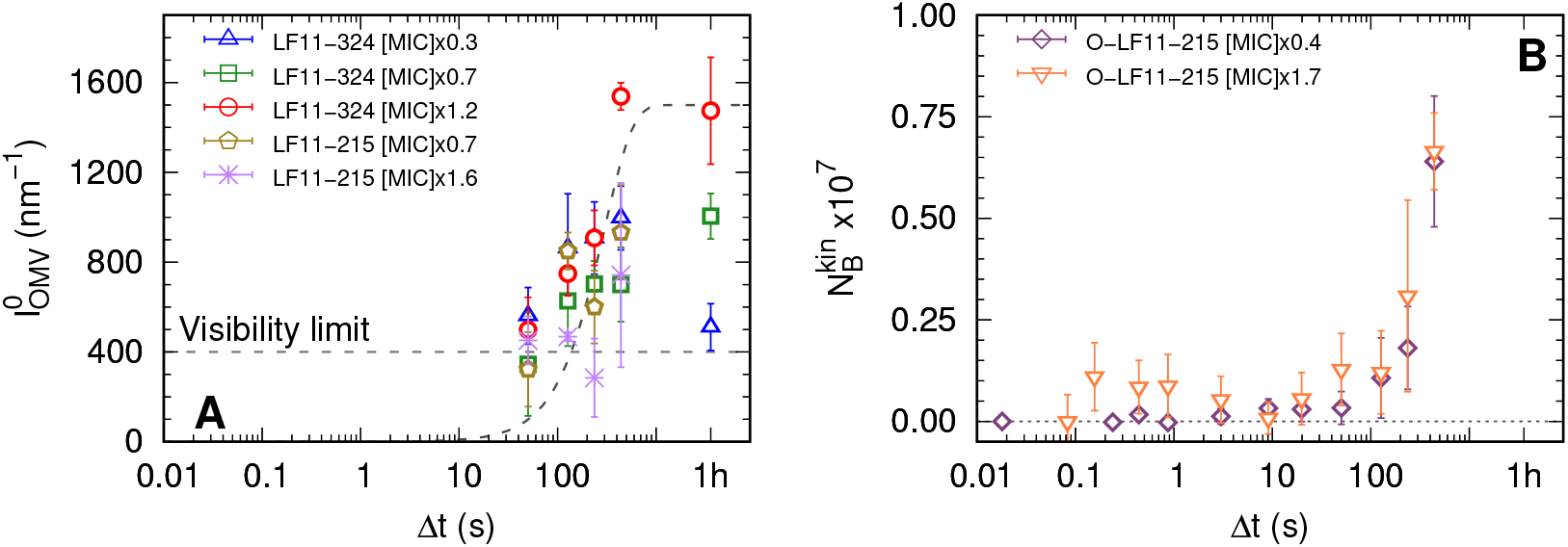
(**A**) Kinetics of the forward intensity of OMV scattering for different concentrations of LF11-324 and LF11-215. The dashed horizontal line represents the detection/‘visibility’ limit, below which *I*_cell_(*q*) + *I*_OMV_(*q*) ≈ *I*cell(*q*) in the entire *q*-range. The dashed exponential curve is a guide for the eyes. (**B**) Evolution of the number of partitioned peptides per cell for two O-LF11-215 concentrations, as calculated from the analysis of *I*_clu_.

### Peptide partitioning and cooperativity

Finally, we applied a previously detailed assay for AMP partitioning in *E. coli* based on growth inhibition (***Marx et al., 2021b***). A statistical analysis of the corresponding data in terms of cumulative distribution functions (see Appendix 4) allowed us to map the probability distributions of inhibiting bacterial growth as a function of cell concentration *n*_cell_, including the minimum concentrations for inhibiting a given percentage *x* of *E. coli*, IC_*x*_ (**Figure 4**); note that MIC ≡ IC_99.9_.

**Figure 4.**
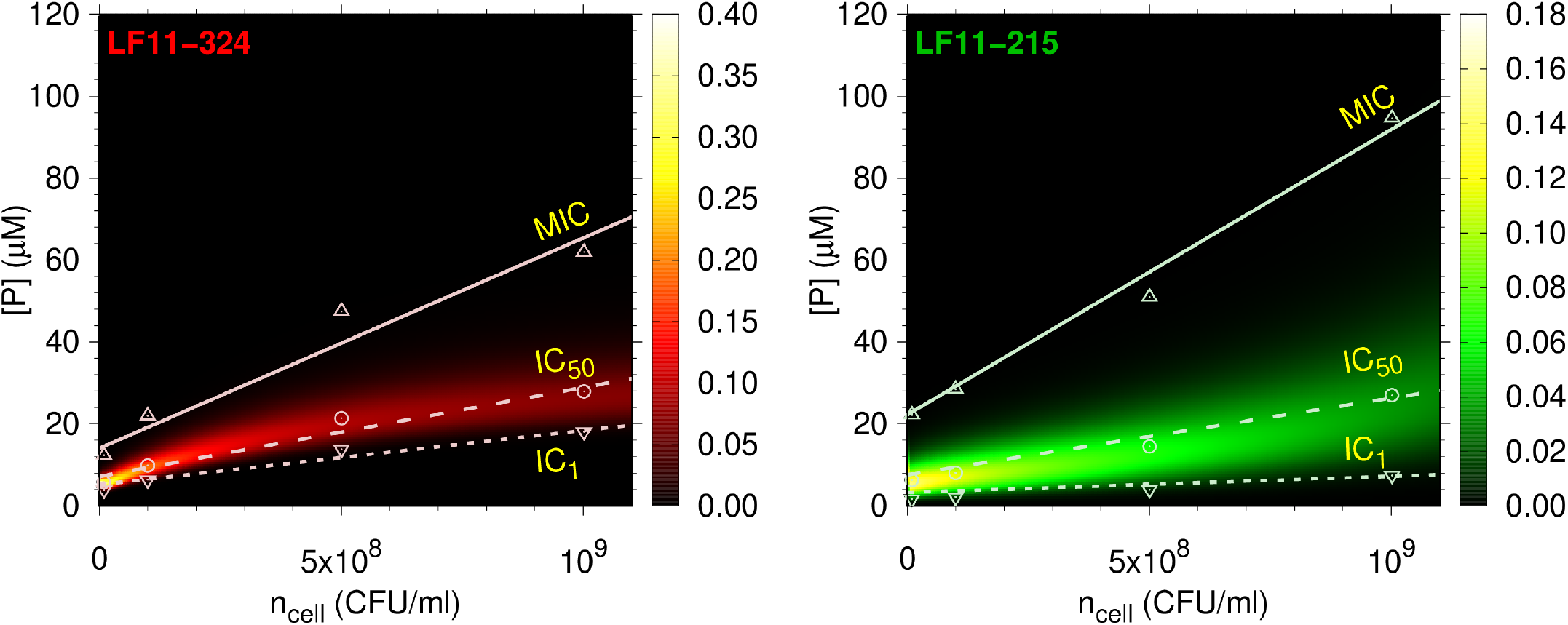
Amount of LF11-324 or LF11-215 required to attain growth inhibited fractions of either 0.999 (MIC, up triangles), 0.5 (circles) or 0.01 (down triangles) in *E. coli* ATCC 25922 as a function of *n*_cell_. Lines are fits with **Equation 1**. These data are overlayed with a surface plot of the associated killing probability density function. The color scales indicate the corresponding magnitudes. **Figure 4–Figure supplement 1.** IC_*x*_ as a function of *ϕ*_IG_ (inverse CDF).

In agreement with our previous report for *E. coli* K12 (***Marx et al., 2021b***), the MIC of LF11-324 is lower than that of LF11-215 at all cell concentrations. For O-LF11-215, MIC values matched those of LF11-324 at low cell concentrations, but increased strongly with *n*_cell_, finally superseding that of LF11-215 and becoming immeasurably high due to peptide aggregation (**Figure 1–Figure Supplement 1**). **Figure 4** additionally illustrates the growth inhibition probabilities (see Appendix 4), whose peaks are close to IC_50_. The distributions are much sharper for LF11-324 than for LF11-215, suggesting an increased cooperativity of killing for LF11-324. The broadness of the LF11-215 killing probability distributions instead caused an earlier onset of growth inhibition at low *n*_cell_. Further, the full width at half maximum of the probability distributions, *σ*_[P]_, increased with cell concentration, e.g. from *σ*_[P]_ ≃ 2.6 *μ*M at *n*_cell_ = 10^7^ CFU/ml to *σ*_[P]_ ≃ 13 *μ*M at *n*_cell_ = 10^9^ CFU/ml for LF11-324. Significant noise levels in growth inhibition data for O-LF11-215 impeded a determination of killing probabilities at inhibitory concentrations < IC_50_. However, data retrieved at higher inhibitory concentrations suggest that the probability distributions roughly match those of LF11-324 at low cell concentrations, but definitely become broader than that of LF11-215 at high cell content (**Figure 4–Figure Supplement 1**). This is another signature of loss of killing efficacy at high *n*_cell_, most likely due to peptide self-aggregation as discussed above.

Next, we derived for each IC_*x*_ (or inhibited fraction, *ϕ*_IG_) the number of cell-partitioned peptides per cell, *N_B_*, and the effective partitioning coefficient, *K^eff^*, applying a previously reported thermodynamic formalism [(***Marx et al., 2021b***); see also **Equation 1**]. *N_B_* increased for all three peptides with *ϕ*_IG_, although the changes of *N_B_* were smallest for LF11-324, followed by LF11-215 and O-LF11-215 **Figure 5**A. These results were confirmed independently also by Trp-fluorescence spectroscopy (see Appendix 3).

**Figure 5.**
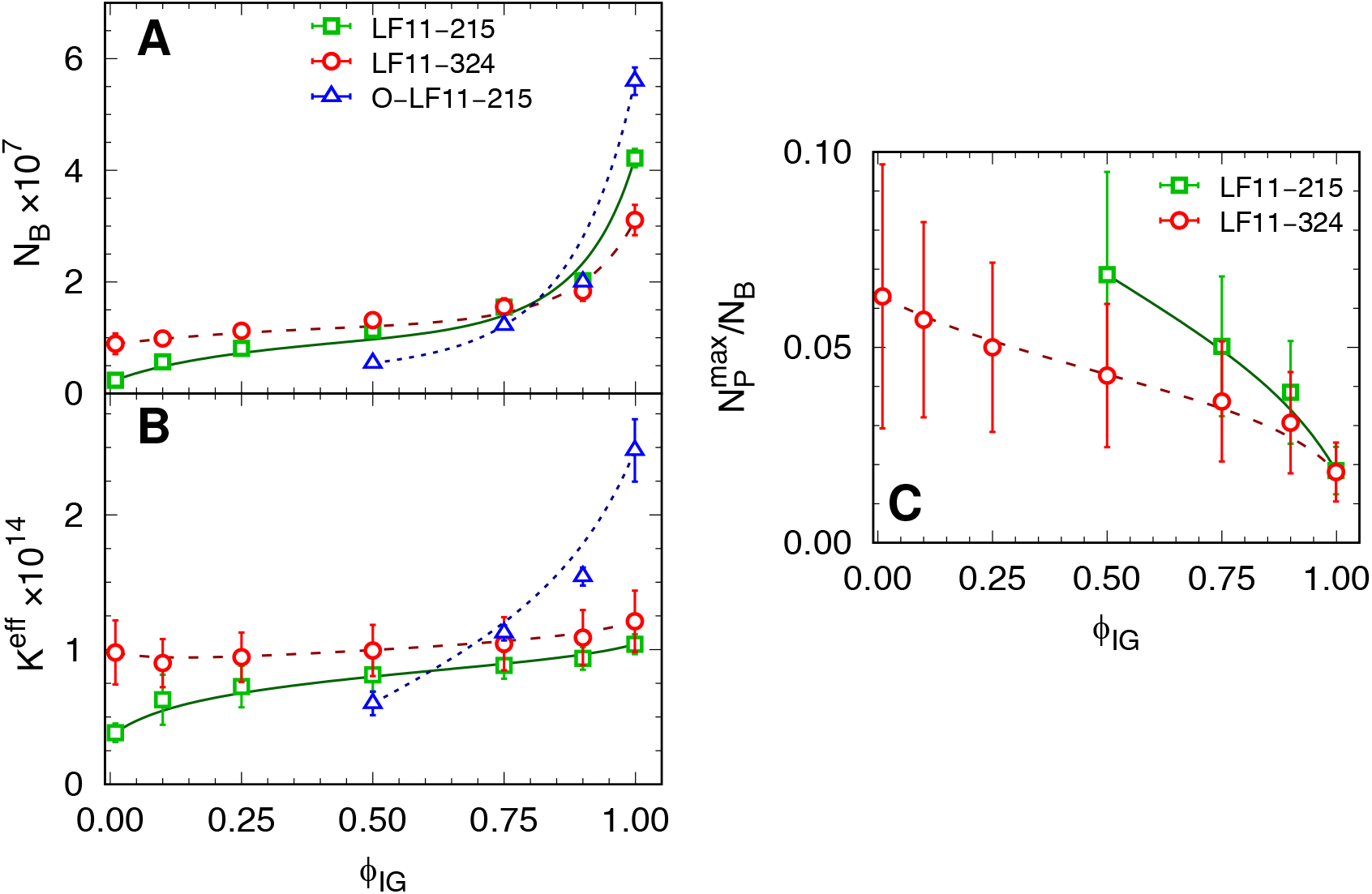
(**A-B**) *N_B_* and *K^eff^* values as a function of inhibited fraction. In the case of O-LF11-215, 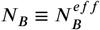. (**C**) Ratio between the maximum number of peptides on the outer leaflet and total number of partitioned peptides, 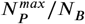, as a function of inhibited fraction. Lines are guides for the eye. **Figure 5–Figure supplement 1.** *ζ*-potential and size measurements of LF11-215 and LF11-324. **Figure 5–Figure supplement 2.** *ζ*-potential and size measurements of O-LF11-215.

This behaviour was also mirrored in the *ϕ*_IG_-dependence of *K^eff^*, which was nearly constant for LF11-324, increased only slightly for LF11-215 and showed the largest variation for O-LF11-215, reaching about 2.5 times higher levels than the other two peptides **Figure 5**B. The approximate equal *K^eff^* values of LF11-324 and LF11-215 for *ϕ*_IG_ > 0.5 demonstrate that both peptides partition about equally well into *E. coli*, not only at the MIC, but in a wide range of *ϕ*_IG_ values.

*ζ*-potential measurements helped to further differentiate between the activity of the two peptides, by determining an upper estimate for the maximum number of peptides associated to the LPS leaflet, 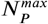 (***Marx et al., 2021b***). After an initial increase of *ζ/ζ*_0_ at low peptide concentrations, plateau-values were rapidly reached for [*P*] ⪆ 0.3×MIC at *ζ/ζ*_0_ = 0.80 ± 0.16 for LF11-215 and *ζ/ζ*_0_ = 0.85 ± 0.17 for LF11-324, where *ζ*_0_ refers to the reference system, i.e. neat bacteria (**Figure 5–Figure Supplement 1**). This converts to 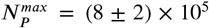 and (6 ± 2) × 10^5^ for LF11-215 and LF11-324, respectively, and to the *ϕ*_IG_-dependence of 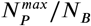 shown in **Figure 5**C. Strikingly, this demonstrates that ~ 98% of the peptides are located within the inner compartments of *E. coli* at the MIC. The fraction of outer-leaflet-partitioned peptides increased toward lower *ϕ*_IG_, and some-what stronger for LF11-215, but does not exceed 10%. An analogous analysis for O-LF11-215 was impeded by the peptide aggregates, whose sizes were on the same order or even larger than that of bacteria (**Figure 5–Figure Supplement 2**).

## Discussion

In agreement with previous studies (***Zweytick et al., 2011, 2014***) we found that end-states of *E. coli* after treatment with either peptide are comparable in terms of cell-wall damage, despite significantly different MICs. Even peptide concentrations far below the MIC led to similar structural effects, including, e.g., loss of positional correlations of the inner and outer membrane, or OMV formation. O-LF11-215, as opposed to LF11-215 and LF11-324, additionally lead to tubulation from the outer membrane (**Figure 1**).

Except for *R*, *ρ*_CP_ and *ρ*_PP_ the kinetic pathways of structural events are equivalent for LF11-215 and LF11-324. Also O-LF11-215 caused comparable variations of the above mentioned parameters on similar time scales. In this case a fully detailed analysis, however, is challenged by the propensity of O-LF11-215 to aggregate in buffer solution. We will thus focus primarily on LF11-215 and LF11-324.

**Figure 6** summarizes the major findings of the present study. The attack of AMPs first shows up in our time-resolved data by changes of the LPS packing density, as well as periplasmic and cytoplasmic SLDs. The decrease of *ρ*_CP_ is associated to a loss of small molecules (<1 kDa)—the major contributors to its X-ray SLD (***Semeraro et al., 2021***)—from the cytoplasm. These molecules first diffuse into the periplasm and then further into extracellular space. The combination of these effects leads to a net increase of *ρ*_PP_. Note that outward net leakage of cytosolic material follows from the fact that final *ρ*_PP_ levels do not reach those of *ρ*_CP_, despite the much larger cytosolic volume. Further, the initial SLDs of buffer and periplasm are comparable, also explaining why our technique is not directly detecting outer membrane leakage. Hence, either observed change of *ρ*_PP_ or *ρ*_CP_ is due to a permeabilization of both cytoplasmic and outer membranes. For LF11-324, the permeabilization of the cytoplasmic membrane occurred as fast as 3 – 10 s after mixing at [P] = 1.2×MIC. Dropping peptide concentration led to a slowing down of this effect (10 – 20 s for [P] = 0.7×MIC, and 50 – 100 s for [P] = 0.3×MIC, **Figure 2**). AMPs need to translocate all the way through the cell wall in order to induce such effects. Considering that resealing of inner and outer membranes potentially cause a delayed onset of leakage consequently implies that peptide translocation possibly proceeds on time scales faster than Δ*ρ*_PP_ or Δ*ρ*_CP_. The drop of *R* is a natural consequence of the loss of cellular content, but occurs at somewhat later times. This is most likely due to the stored elastic energies of the peptidoglycan layer, which will initially resist rapid deformations (***Yang et al., 2018***). Note, however, that the peptidoglycan properties are likely affected by direct interactions with peptides (***Zhu et al., 2019***). Remarkably, cytoplasmic membrane leakage occurs up to ~ 30 times later for LF11-215, independent of peptide concentration (**Figure 2–Figure Supplement 3**).

**Figure 6.**
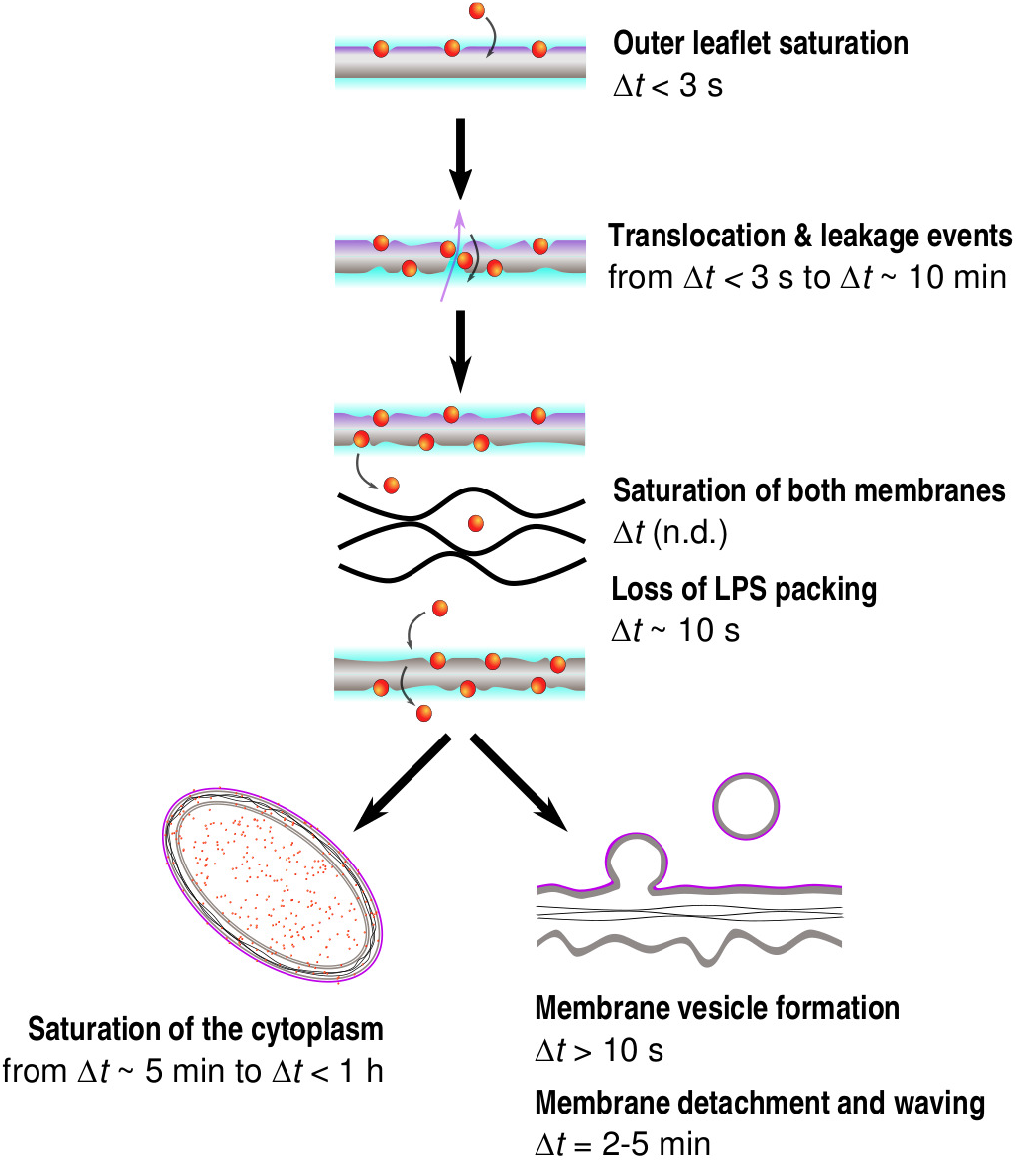
Simplified time sequence of LF11-215 and LF11-324 mode-of-action. The outer leaflet saturates with peptides within the first seconds after their attack. Then, depending of AMP type and concentration, a number of rare translocation events, coupled with leakage, takes places over a broad time range. When both membranes are saturated with peptides (exact time not determined), the cell-wall breaks down, leading to OMV formation (Δ*t* >10 s), detachment of outer and inner membranes and waving (Δ*t* >2–5 min). Simultaneously, AMPs saturate the internal compartments within several minutes, again depending of AMP type and concentration.

Pronounced differences between LF11-324 and LF11-215 were also observed from their efficacies as a function of cell concentration (**Figure 4**). At equal *n*_cell_, growth-inhibition probability distributions are much narrower for LF11-324. Apparently, this increased ‘cooperativity’ correlates with the peptide’s ability to permeabilize the cytoplasmic membrane faster. It is further illuminating to discuss the total amount of peptide penetrating into the cytosol. Both peptides saturate the outer LPS leaflet already at concentrations lower than 0.3×MIC (**Figure 5–Figure Supplement 1**), corresponding to an ~ 1:3 AMP/LPS molar ratio as upper boundary estimate. Thus peptides penetrate the outer membrane already when effects on bacterial growth are still very small. Assuming that LF11-324 and LF11-215 associate to first order at comparable amounts with other membrane leaflets leads to a lower boundary estimate of ~ 92% of all pepides being located in the cytosol at the respective MICs. This corresponds to huge cytosolic concentrations of ~ 80 mM for LF11-324, and ~ 110 mM for LF11-215. The difference in cytosolic concentrations between the two peptides corresponds to about the difference in MICs (**Figure 1–Figure Supplement 1**). Our finding that only a minor fraction of the peptides are located within the cell wall even persists at sub-MIC concentrations. This explains why the average damage of its structure is not peptide specific and does not depend on peptide concentration.

Wimley discussed already about 10 years ago the consequences of having 10^7^ to 10^9^ AMPs per cell (***Wimley, 2010***), suggesting a ‘reservoir’ of non-membrane-bound peptides that would outnumber proteins, ATP and other metabolites. Cytosolic targets were also confirmed by our TEM data [and those of others (***Hammer et al., 2010***; ***Scheenstra et al., 2019***)] showing a collapsed nucleoid region (**Figure 1**). For instance this could be due to a competition mechanism with polyamines, as putative stabilizers of the functional architecture of the DNA ring (***Hou et al., 2001***). In support of the hypothesis of cytosolic targets, recent solid-state ^31^P-NMR measurements of *E. coli* in the presence of AMPs revealed increased dynamics of DNA and RNA phosphate groups correlated to TEM observations of collapsed nucleoid volume (***Overall et al., 2019***). In addition, however, AMP interactions with other negatively charged metabolites and macromolecules in the cytosol, such as ribosomes and proteins, should be considered as potentially detrimental to the bacteria (***Zhu et al., 2019***). The about 1.4 times lower cytosolic concentration of LF11-324 then is a signature of a higher potency as compared to LF11-215 to interfere with one or more of the above listed candidates, hampering a number of metabolic functions (***Scocchi et al., 2015***), or inducing an apoptosis-like mechanism (***Dwyer et al., 2012***).

It follows from the considerations above that bacteria have an increased probability to recover from the peptides’attack, if the cytosolic concentrations of LF11-324 and LF11-215 fall below those reported above (i.e. ⪅ 80 and ⪅ 110 mM, respectively). Thus, both the ability of LF11-324 to swiftly translocate through the cell envelope and its higher propensity to interfere with the metabolic machinery contribute to its higher cooperativity in killing *E. coli* (**Figure 4**). Our experimental setup is not sensitive to directly observe either transient membrane pores or other membrane defects. Note, however, that previous studies in lipid-only mimics of bacterial membranes showed only weak membrane remodeling effects of lactoferricin derivatives as compared to other peptides (***Zweytick et al., 2011***; ***Marx et al., 2021b***). This might be also a consequence of the rather short amino acid sequence of the lactoferricin derivatives, implying a too small length (~ 1 nm) to span the whole membrane thickness (3 – 4 nm). In turn, the highly flexible secondary structure and an amphipathic momentum being aligned along the peptide’s backbone (***Zorko et al., 2009***; ***Zweytick et al., 2011, 2014***) should allow LF11-324 and LF11-215 to translocate membranes at higher rates than observed for linear peptides (***Ulmschneider, 2017***; ***Kabelka and Vácha, 2018***). Note that peptide translocation can also lead to transient membrane leakage events (***Ulmschneider, 2017***), even with negligible AMP-induced lipid flip-flop (***Marx et al., 2021a***). The higher hydrophobicity of O-LF11-215 should increase the likelihood of remaining membrane bound, which might build up differential membrane curvature stress and lead to the observed formation of membrane tubules. We also note that the different leakage kinetics for the LF11 peptides suggest a coupling to translocation kinetics, which in turn depends on membrane partitioningof the AMPs. Indeed, recently reported data for cytoplasmic membrane mimics of cardiolipin, phosphatidylethanolamine and phosphatidylglycerol show a somewhat faster membrane partitioning for LF11-324 than for LF11-215 (***Marx et al., 2021b***).

Concluding, the superior time-resolution and sensitivity to structural changes from cellular size to molecular packing of synchrotron USAXS/SAXS allowed us to demonstrate, upon combination with advanced data modeling and complementary techniques, that AMPs are able to reach the cytosolic compartment of bacteria on the seconds time scale and thus much faster than previously considered (***Sochacki et al., 2011***). Two factors emerge as key components for AMP efficacy: (*i*) a fast translocation through inner and outer membranes, rapidly reaching extremely high cytosolic AMP concentration levels (~ 100 mM), and (*ii*) an efficient shut-down of the bacterial metabolism, i.e., the lower the number of ‘needed’ AMPs in the cytosol the better. This latter process is driven by interactions of the AMPs with yet-to-be-determined cytosolic molecules, but most likely candidates are the polyanionic DNA, RNA, ribosomes and proteins (***Zhu et al., 2019***), or charged metabolites. Collateral damage of the cell-wall structure is a ‘by-product’ of AMP activity. That is, it occurs already at sub-MIC concentrations (due to a rapid saturation of membranes with peptides) and thus is not the primary cause for growth inhibition. It is currently not clear whether the present findings can also be extended to other AMPs. Yet, similar conclusions were drawn for the peptide LL-37 by ***Zhu et al.*** (***2019***), and an emergent number of AMPs has been reported to show comparable partitioning behavior in bacteria (***Loffredo et al., 2021***). We thus propose, that the combination of membrane translocation speed and efficient shut down of bacterial metabolism are generic factors that should be considered in designing future AMPs to combat infectious diseases. This also implies a widening of the pure focus on membrane-activity of AMPs currently applied in many studies.

## Methods and Materials

### Samples

*Escherichia coli* ATCC 25922 were provided by the American Type Culture Collection (Manassas, VA). Freeze-dried peptides powder of LF11-215 (H-FWRIRIRR-NH_2_), LF11-324 (H-PFFWRIRIRR-NH_2_) and O-LF11-215 (octanoyl-FWRIRIRR-NH_2_), purity >95%, were purchased from the Polypeptide Laboratories (San Diego, CA). Lysogeny broth (LB)-agar and LB medium were obtained from Carl Roth, (Karlsruhe, Germany). All the other chemicals were purchased from Sigma-Aldrich (Vienna, Austria).

#### Bacterial cultures

Bacterial colonies of *E. coli* ATCC 25922 were grown in LB-agar plates at 37 °C. Overnight cultures (ONCs) were prepared by inoculating a single colony in 3 ml LB-medium in sterile polypropylene conical tubes (15 ml), allowing for growth under aerobic conditions for 12–16 hours in a shaking incubator at 37 °C. Main cultures (MCs) were then prepared by diluting ONCs in 10 ml LB-medium in 50 ml sterile polypropylene conical tubes. Bacteria in the MCs grew under the same conditions applied to ONCs up to the middle of the exponential growth phase. Cells were then immediately washed twice and re-suspended in nutrient-free and isotonic phosphate-buffered saline (PBS) solution (phosphate buffer 20 mM, NaCl 130 mM) at pH 7.4.

#### AMP samples

LF11-324 and O-LF11-215 peptides displayed a weak solubility in PBS. AMP stocks (including LF11-215) were then prepared by adding acetic acid and DMSO, up to 0.3% and 3% vol/vol, respectively. Peptide stock solutions were diluted for measurements yielding a final concentration of 0.01% acetic acid and 0.1% vol/vol dimethyl sulfoxide (DMSO) (final pH 7.2). Hence, possible effects of DMSO on the cell envelope, as observed for model membrane structures (***Gironi et al., 2020***) can be neglected. Control USAXS/SAXS experiments adding a similar amount DMSO and acetic acid to *E. coli* also showed no discernible effect of the organic solvent (data not shown). Similarly, control experiments were performed to exclude effects on bacterial growth. Stock concentrations were determined by measuring the absorption band of the Trp residues with the spectrophotometer NanoDrop ND-1000 (Thermo Fisher Scientific, Waltham, MA). Peptide stock solutions were stored in silanized glass tubes until use.

### Antimicrobial activity and partitioning modeling

The antimicrobial activity of the AMPs on *E. coli* was tested in the bacterial concentration range of 5× 10^5^ to 10^9^ CFU/ml using a modified susceptibility microdilution assay (***Jorgensen and Ferraro, 2009***). Cell suspensions were incubated at a given AMP concentration for 2 h at 37 °C (control samples were incubated in buffer only). Cell growth was monitored upon addition of double concentrated LB-medium for about 20 h using a Bioscreen C MBR (Oy Growth Curves Ab, Helsinki, Finland).

#### Partitioning modeling

The analysis of the inhibited fraction of cells, *ϕ*_IG_, as a function of the total concentration of peptides, [*P*], enabled the extraction of IC_*x*_ values as a function of *n*_cell_, as detailed in Appendix 4. Results were fitted with the linear partitioning function

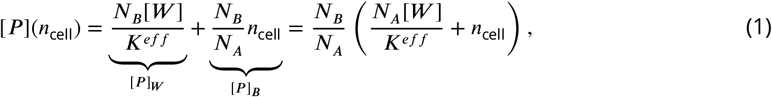

where [*P*]_*W*_ and [*P*]_*B*_ are the concentrations of AMPs dispersed in the aqueous phase and segregated into the cells, respectively; *N_A_* is the Avogadro’s constant; [*W*] is the concentration of water molecules in bulk (55.3 M at 37 °C); *K^eff^* is the effective mole-fraction partitioning coefficient; and *N_B_* is the number of peptide monomers that are partitioned within a single cell [see ***Marx et al.*** (***2021b***) for details].

A similar approach was exploited in the case of O-LF11-215 peptide clusters in solution. In this case, the total peptide concentration is given by [*P*] = [*P*]_*W*_ + [*P*]_*B*_ + *N*[*A*], where [*A*] is the molar concentration of aggregates, each of them consisting of an average number of peptides *N*. We also define the aggregate fraction *f_A_* = *N*[*A*]/[*P*] and assume the equilibrium state *N*[*P*]_*W*_ ⇌ [*A*]. The definition of the molar partitioning coefficient *K_x_* ∝ *K^eff^* (***Marx et al., 2021b***) refers to the balance of concentration of free peptides in bulk and partitioned peptide into the cells. Hence, its bare definition is unaffected by the presence of clusters. Finally, it is trivial to show that in this case **Equation 1** becomes:

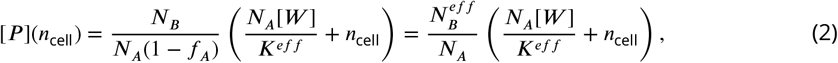

where the fitting parameter 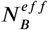 is an upper boundary estimate for the actual number of partitioned peptides per cell 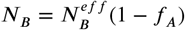.

### *ζ*-potential, cell size, and outer leaflet distribution of peptides

*ζ*-potential and and dynamic light scattering (DLS) measurements were carried out with the Zetasizer Nano ZSP (Malvern Panalytical, Malvern, UK). *E. coli* suspensions were incubated with different concentrations of AMPs in buffer for 1 hour at 37 °C prior to each measurements. Control samples (no AMPs) were suspended and incubated in buffer. A concentration of 10^7^ CFU/ml provides the optimal compromise between high signal-to-noise ratio and low multiple-scattering bias. The AMP concentrations were centered on the MIC values previously determined with the susceptibility microdilution assays, spanning from about 0.2× to 2.5×MIC. For *ζ*-potential measurements the voltage for the electrodes was set to 4 V, such that currents did not exceed 1 mA, because of the high conductivity of the PBS buffer. Further, measurements were paused between repetitions for 180 seconds. This prevented heat productions leading to sample denaturation and accumulation on the electrodes. The experiments were repeated three times and, due to the low sensitivity of such a set-up, each of them consisted of a minimum of six measurements [see also ***Marx et al.*** (***2021b***)]. For each system, *ζ*-potential values and associated errors were given by the medians and the median absolute deviations, respectively, averaging over at least 18 repetitions. The same number of scans was also used to obtain mean and standard deviation values for the hydrodynamic diameter, *d_H_*, of the cells.

From the measured *ζ*-potential values we estimated the maximum number of peptides that partition into the outer LPS leaflet, 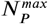, as reported recently (***Marx et al., 2021b***)

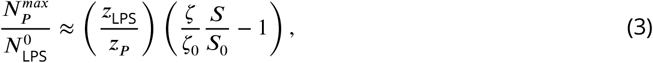

where *z*_LPS_ = −6 (***Wiese et al., 1998***) and *z_P_* = +5 (***Zweytick et al., 2011***) are the nominal charges of LPS and LF11-215 or LF11-324 AMPs, respectively; *ζ* and *S* are the *ζ*-potential and total surface values of the system upon addition of peptides; and *ζ*_0_ and *S*_0_ are the respective reference values (no AMPs). 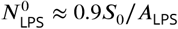 is the estimated number of LPS molecules, where *A*_LPS_ ≃ 1.6 nm^2^ (***Kim et al., 2016***; ***Micciulla et al., 2019***) is the lateral area per LPS molecule. The prefactor originates from considering a maximum surface coverage of 90% by LPS molecules (***Seltmann and Holst, 2002***). *S*_0_ was derived from DLS measurements, approximating the bacterial shape by a cylinder 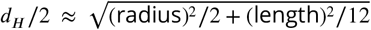, considering that the hydrodynamic radius is approximately equivalent to the radius of gyration for micron-sized objects. Then fixing the radius at about 400 nm (***Semeraro et al., 2021***) and retrieving the length from the above relation for *d_H_* one obtains *S*_0_ ≈ 5 × 10^6^ nm^2^.

### Fluorescence spectroscopy

Fluorescence spectroscopy experiments were done with the Cary Eclipse Fluorescence Spectrophotometer (Varian/Agilent Technologies, Palo Alto, CA). The excitation wavelength was set to *λ* =280 nm (which corresponds to the maximum intensity of the absorption/excitation band of Trp), and emission spectra were acquired in the *λ*-range between 290 and 500 nm, with the Trp emission band peak being expected to lie around 330 to 350 nm. Samples were loaded in quartz cuvettes of 1 cm path-length. The background was subtracted from every Trp-spectrum prior to further analysis.

#### Peptide solubility

Trp emission allowed determining whether LF11 peptides form aggregates or not in the MIC range. Spectra of LF11-only samples at [*P*] =100 *μ*g/ml were fitted with the log-normal-like function (***Burstein and Emelyanenko, 1996***; ***Ladokhin et al., 2000***)

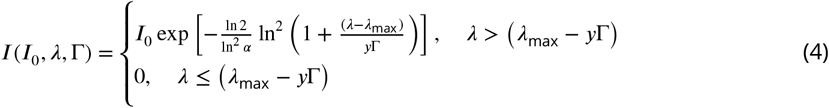

where *λ*_max_ and *I*_0_ are, respectively, wavelength and intensity of the emission peak; Γ is the full-width-at-half-maximum (FWHM)of the band; *α* is a skewness parameter (fixed at an optimum value of 1.36 after testing; and *y* = *α*/(*α*^2^ – 1).

Both LF11-215 and LF11-324 (see Appendix 1) showed a peak at about *λ*_max_ ≃ 353 nm and Γ ~ 63 nm. This is consistent with a location of the Trp residues in polar chemical environments having full mobility and thus suggests that these AMPs are monomeric (***Burstein and Emelyanenko, 1996***). In contrast, the acylated O-LF11-215 showed a significant blue-shift related to a location of Trp within apolar surroundings (***Ladokhin et al., 2000***), indicating the formation of peptide aggregates.

#### Peptide partitioning

Analogously to the partitioning analysis performed for lipid only membranes (***Marx et al., 2021b***), the Trp emission of bacteria/AMPs mixtures were treated as a two-component signal, one coming from the peptides in the aqueous phase, and the second one from AMPs interacting with the cells.

Bacterial suspensions were incubated with different concentrations of AMPs in buffer for 1 hour at 37 °C (incubator Thermomixer C, Eppendorf, Germany). Reference samples (no AMPs) were suspended and also incubated in PBS. Experiments were performed at cell concentrations of 5×10^7^, 10^8^ and 5×10^8^ CFU/ml, and AMPs amounts equal to 0.5×, 1× and 2×MIC (each experiment was repeated three times). Fluorescence intensities were background-subtracted using the bacteria-only reference spectra at the corresponding concentrations. This enabled us to subtract the average signal from the aromatic residues in the cells, and the scattering arising from the high concentration of the cell suspensions. Spectra were analyzed with a linear combination of two independent bands (see **Equation 4**) *I^W^* and *I^B^*, referring to AMPs in bulk (W) and partitioned into the lipid bilayer (B). *λ^W^* and Γ^*W*^ were fixed to the reference values obtained by analyzing spectra from pure AMPs, and 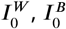, *λ^B^* and Γ^*B*^ were freely adjusted. LF11-only solutions were measured to calibrate their intensity dependence in buffer. Then, the retrieved 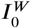 values were converted to the concentration of dissociated peptides [*P*]_*W*_. This allowed us to obtain the so-called peptide bound-fraction as *f_B_* = 1 – [*P*]_*W*_/[*P*]. The aggregation of O-LF11-215 led to low 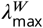 values (see Appendix 1) and precluded the a similar analysis for these peptides.

### Transmission electron microscopy

*E. coli* suspensions at 5×10^8^ CFU/ml were mixed with peptides at the corresponding MICs and 0.5×MICs in buffer A, and incubated for 1 hour at 37 °C (incubator Thermomixer C, Eppendorf, Germany). Control samples (no AMPs) were suspended and incubated in buffer A. Cell suspensions were centrifuged at 1300 *g* for 4 minutes in 1.5 ml Eppendorf tubes and resuspended (fixed) in 3% vol/vol glutaraldehyde and 0.1 M cacodylate buffer to bring the rapid cessation of biological activity and to preserve the structure of the cell. After fixation the samples were washed and post-fixed in 1% vol/vol OsO_4_ in 0.1 M cacodylate buffer. Dehydration was carried out in an ascending ethanol series followed by two steps with propylene oxide (***Hayat, 1989***) and embedded in Agar Low Viscosity Resin (Agar Scientific, Stansted, UK). Ultrathin sections (70–80 nm) were prepared on an Ultramicrotome UC6 (Leica Microsystems, Vienna, Austria) equipped with a 35° Ultra Diamond-knife (Diatome, Nidau, Switzerland). The grids were poststained with Uranyless (Science Services, Munich, Germany) and lead citrate according to Reynolds (***Hayat, 1989***). Transmission electron microscopy images were acquired with Tecnai T12 at 120kV (TFS, Warmond, Netherlands).

### Small angle scattering

#### (Ultra-) Small-angle X-ray scattering

USAXS/SAXS measurements were performed on the TRUSAXS beamline (ID02) at the European Synchrotron Research Facility (ESRF), Grenoble, France. The instrument uses a monochromatic beam (*λ* =0.0995 nm) that is collimated in a pinhole configuration. Measurements were performed with sample-to-detector distances of 30.8 and 3.0 m, covering a *q*-range of 0.001-2.5 nm^−1^ (***Narayanan et al., 2018***). Two-dimensional scattering patterns were acquired on a Rayonix MX170 detector, normalized to absolute scale and azimuthally averaged to obtain the corresponding one-dimensional USAXS/SAXS profiles. The normalized cumulative background from the buffer, sample cell and instrument were subtracted to obtain the final *I*(*q*). Bacterial samples (concentration ~ 10^9^ CFU/ml) were incubated with peptides for one hour at 37 °C and directly measured in a quartz capillaries of 2 mm diameter (37 °C), mounted on a flow-through set-up in order to maximize the precision of the background subtraction. Time-resolved experiments (*n_cell_* ~ 10^9^ CFU/ml) were instead performed with a stopped-flow rapid mixing device (SFM-3/4 Biologic, Seyssinet-Pariset, France), with 50 ms mixing of bacterial and peptides stock suspensions (37 °C), and enabling data acquisition after a kinetic time of about 2.5 ms (***Narayanan et al., 2014***). A total of 50 frames was recorded for each experiment with an exposure time of 0.05 seconds and a logarithmic time-spacing ranging from 17.5 ms to about 10 minutes. Radiation damage tests were performed on reference systems prior to setting this X-ray exposure-times. The scattering intensities were further corrected for sedimentation and background scattering from the stopped-flow cell.

#### Contrast-variation small angle neutron scattering

SANS experiments were performed on the D11 instrument at the Institut Laue-Langevin (ILL), Grenoble, France, with a multiwire ^3^He detector of 256 × 256 pixels (3.75 × 3.75 mm^2^). Four different set-ups (sample-to-detector distances of 2, 8, 20.5, and 39 m with corresponding collimations of 5.5, 8, 20.5 and 40.5 m), at a wavelength *λ* =0.56 nm (Δ*λ*/*λ* =9%), covered a *q*-range of 0.014–3 nm^−1^. Two distinct *E. coli* suspensions were incubated with peptides LF11-215 and LF11-324 in buffer for one hour at 37 °C. The bacterial concentration during the incubation was 10^9^ CFU/ml, and both peptide concentrations were in the range of the measured MICs. Samples were then washed and resuspended in five different PBS solutions, containing 10, 30, 40, 50, or 90 wt% D_2_O. Samples (concentration ~ 10^10^ CFU/ml) were contained in quartz Hellma 120-QS banjo-shaped cuvettes of 2 mm pathway and measured at 37 °C. Cuvettes were mounted on a rotating sample holder, which prevented the bacteria from sedimenting. Data were reduced with the Lamp program from ILL, performing flat field, solid angle, dead time and transmission correction. Further data were normalized by the incident neutron flux (via a monitor), and corrected by the contribution from an empty cell. Experimental set-up information and data are available at https://doi.ill.fr/10.5291/ILL-DATA.8-03-910.

Note that the present experimental time (~ 2 h) is much shorter than the onset of cell lysis (***Zweytick et al., 2011***). As control, SANS measurements were repeated at extended times after mixing with the AMP in order to test for sample stability (data not shown), in terms of shape, cell-wall structure and densities. The scattering intensities after 2, 4, 6 and 8 hours were comparable (with the exception of a weak decrease of *ρ*_CP_ between 2 and 4 hours). Further a comparison with TEM and SAXS data suggests that the peptide-induced cell-damage is stabilized within one hour, and does not develop any further for at least 8 hours.

#### Data analysis: peptide clusters

Reference O-LF11-215 samples were measured in the MIC range to investigate the microstructure of the peptide clusters. The SAXS pattern of O-LF11-215 was fitted with the equation

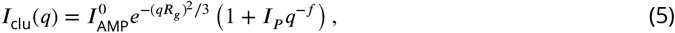

where the term in brackets is related to the structure of the aggregates, and the exponential Guinier function accounts for the form factor of their subunits of radius of gyration *R_g_* (***Zemb and Lindner, 2002***), and 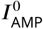 is forward scattering intensity. The function *I_P_q^−f^* is the Porod law that describes the high-*q* asymptotic trend of scattering signal from the aggregates (***Glatter et al., 1982***), where *I_P_* is a scaling factor that depends on the surface properties of the aggregates, and *f* is related to their fractal dimension (***Sorensen, 2001***) (see Appendix 1).

#### Data analysis: bacterial modeling

X-ray and neutron scattering data were jointly analyzed with a recently reported analytical scattering model (***Semeraro et al., 2021***). USAXS/SAXS patterns of end-states displayed an excess scattering contribution between *q* ~ 0.1–0.2 nm^−1^ in the case of LF11-215 and LF11-324, not visible in the corresponding SANS patterns. Note that SANS experiments were conducted on samples that were washed and resuspended in different D_2_O-containing buffer, while SAXS data were acquired immediately after one hour incubation with peptides. Together with the observation of OMVs by TEM (**Figure 1**), this suggests that the additional scattering contribution in SAXS data could be due to freely diffusing OMVs in the suspension medium.

All scattering data were fitted with the analytical functions

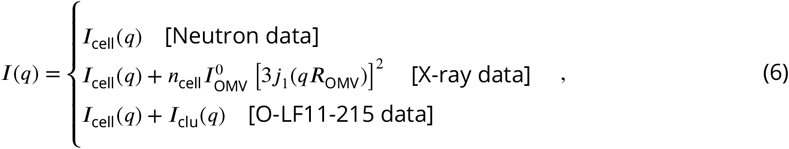

where *I*_cell_(*q*) is the scattering form factor for *E. coli*, as reported in ***Semeraro et al.*** (***2021***), and 3*j*_1_(*qR*_OMV_) is the form factor of a sphere of radius *R*_OMV_, with *j*_1_ being the normalized spherical Bessel function of order 1. The prefactor 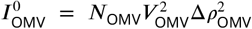 is the OMV forward scattering, where *N*_OMV_, *V*_OMV_ and Δ*ρ*_OMV_ are, respectively, their number, volume and SLD difference to the buffer. The choice of a simple spherical form factor was driven by its simplicity for checking whether bacteria and OMVs have to be considered as non-interacting scatterers or not. Tests using an interaction cross-term approximating budding OMVs by spheres decorating a larger surface (***Larson-Smith et al., 2010***), did not result in significant contributions, confirming the dominance of freely diffusing OMVs. Note also that the 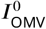 values were independent of the shape of the normalized form factor, as they include our estimation of *V*_OMV_ and Δ*ρ*_OMV_ (Appendix 2). Finally, in the case of SANS data, instrumental smearing was taken into account. Data were fitted with a convolution of *I*(*q*) and a Gaussian function with *q*-dependent width values, as provided by the reduction tools at D11. In USAXS/SAXS data, the smearing effect was negligible.

After thorough testing, the analysis of SAS data (end-states and kinetics) was conducted using only seven adjustable parameters describing *I*_cell_(*q*). These were the number of LPS molecules, *N*_OS_; the cytoplasm radius, *R*; the scattering length densities (SLDs) of the cytoplasmic, *ρ*_CP_, and perisplamic space, *ρ*_PP_; the periplasmic average thickness, Δ_OM_, and its deviation, *σ*_OM_; and the SLD of the peptidoglycan layer, *ρ*_PG_. Additionally 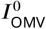 and *R*_OMV_ were fitted for scattering intensities in the presence of LF11-215 and LF11-324, while 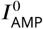 was used and adjusted in the case of O-LF11-215. Other parameters of *I*_clu_(*q*) were fixed according to the O-LF11-215 alone systems (see **Table 1**). This allowed us to fully describe the scattering-pattern variations upon peptide activity. Other parameters, including those accounting for the structure of inner and outer membranes, were fixed to the references values [see complete description in ***Semeraro et al.*** (***2021***); all fixed parameters are listed in **Table 2** and **Table 3**.

**Table 2.**
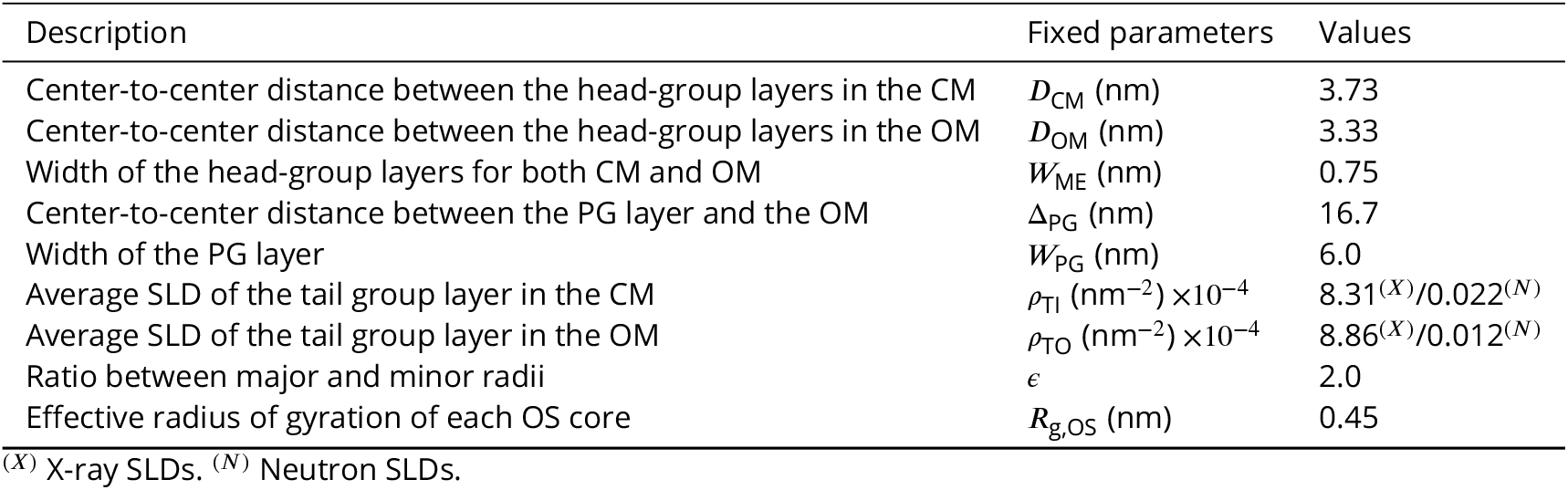
List of fixed parameters for the combined analysis of USAXS/SAXS and contrast variation SANS data of *E. coli*.

| Description | Fixed parameters | Values |
| --- | --- | --- |
| Center-to-center distance between the head-group layers in the CM | $D_{CM}$ (nm) | 3.73 |
| Center-to-center distance between the head-group layers in the OM | $D_{OM}$ (nm) | 3.33 |
| Width of the head-group layers for both CM and OM | $W_{ME}$ (nm) | 0.75 |
| Center-to-center distance between the PG layer and the OM | $\Delta_{PG}$ (nm) | 16.7 |
| Width of the PG layer | $W_{PG}$ (nm) | 6.0 |
| Average SLD of the tail group layer in the CM | $\rho_{TI}$ (nm<sup>-2</sup>) $\times 10^{-4}$ | $8.31^{(X)}/0.022^{(N)}$ |
| Average SLD of the tail group layer in the OM | $\rho_{TO}$ (nm<sup>-2</sup>) $\times 10^{-4}$ | $8.86^{(X)}/0.012^{(N)}$ |
| Ratio between major and minor radii | $\epsilon$ | 2.0 |
| Effective radius of gyration of each OS core | $R_{g,OS}$ (nm) | 0.45 |

<sup>(X)</sup> X-ray SLDs. <sup>(N)</sup> Neutron SLDs.

**Table 3.**
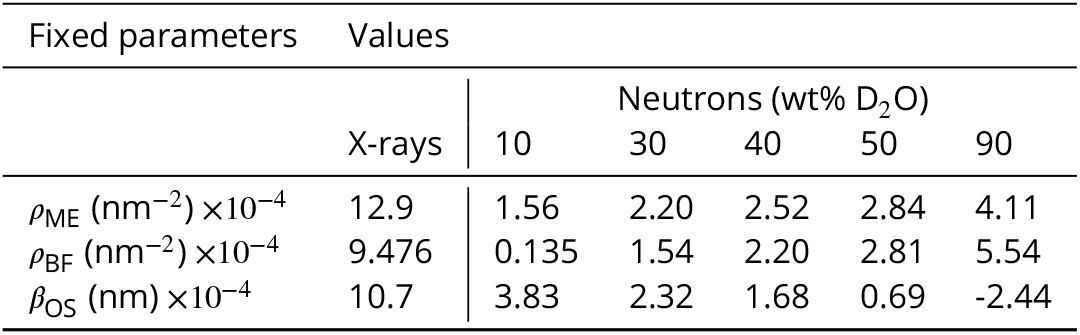
List of fixed and D_2_O-dependent parameters for the combined analysis of USAXS/SAXS and contrast variation SANS data of *E. coli*. The average SLD of both CM and OM head-group layers, *ρ*_ME_, the SLD of the buffer solution, *ρ*_BF_, and the product of the each OS core volume and it contrast relative to the buffer, *β*_OS_ = *V*_OS_Δ*ρ*_OS_.

The scattering intensities of O-LF11-215-aggregates was comparable to that of bacteria in the high *q*-range (**Figure 2–Figure Supplement 1**D). While this affected the quality of the ultrastructural parameters, it enabled at the same time the investigation of the kinetics of the AMP uptake. Indeed, by assuming that O-LF11-215 is primarily forming aggregates in solution, 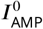 at Δ*t* = 17.5 ms can be converted to the total known peptide concentration [*P*]. Hence, the further assumption that peptides leaving the clusters are directly partitioning into the cell allows to convert the difference 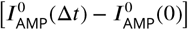 to [*P*]_*B*_ (Δ*t*). It follows that

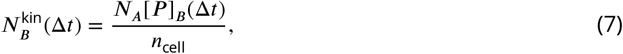

where 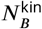 is the number of O-LF11-215 partitioned within the volume of a single cell that can be obtained time-resolved USAXS/SAXS data.

Finally, time-resolved USAXS/SAXS data were fitted using the parameters of the initial [see ***Semeraro et al.*** (***2021***)] and end-states as boundaries and guide to refine the *χ*^2^ minimization. This was accomplished by means of a genetic selection algorithm exploiting >300 repetitions of converging fittings [see details in ***Semeraro et al.*** (***2021***)]. Mean values and errors of the adjustable parameters from both USAXS/SAXS and SANS data are the mean and standard deviation values of the *>*300 converging series. Variations in Δ_OM_ and *σ*_OM_ at Δ*t* = 17.5 ms are due to lower signal-to-noise ratio available in time-resolved measurements. Compared to the reference system, stopped-flow measurements were performed with a lower exposure time and *E. coli* concentration [see ***Semeraro et al.*** (***2021***)].

## Acknowledgments

This project was supported by the Austrian Science Funds (FWF), grant no. P 30921. ESRF – The European Synchrotron and the Institut Laue-Langevin (ILL) are acknowledged for provision of SAXS (proposals LS-2513 and LS-2869) and SANS (exp. 8-03-910) beamtimes. The authors are grateful to T. Narayanan for his invaluable support, and thank J. M. Devos, D. Marquardt and M. Pachler, for their support during the proof-of-concept experiments (LS-2513), and the biological support laboratory at EMBL Grenoble for providing access to the laboratory equipment for bacterial sample preparation. The authors also acknowledge N. Malanovic for sharing her expertise about bacterial cultures, and to S. Keller for the fruitful discussions. Finally, the authors thank the whole staff of ID02 and the D11 for support and availability.

## Appendix 1 Clusters of acylated peptide O-LF11-215

Peptide clusters formed by O-LF11-215 were investigated by Trp fluorescence and USAXS/SAXS. Trp spectra displayed a *λ*_max_ ≃ 336 nm and a FWHM of about 67 nm, which can be related to a heterogeneous distribution of Trp with apolar surroundings (***Ladokhin et al., 2000***). In addition, O-LF11-215 exhibited Trp phosphorescence emission at about 450 nm, which usually is not measurable due to its dynamic quenching by oxygen and impurities in aqueous suspensions (***Cioni and Strambini, 2002***). Its presence suggests that a significant portion of Trp residues are buried in hydrophobic cores, with no access to the solvent and with a local high viscosity.

**Appendix 1 Figure 1.**
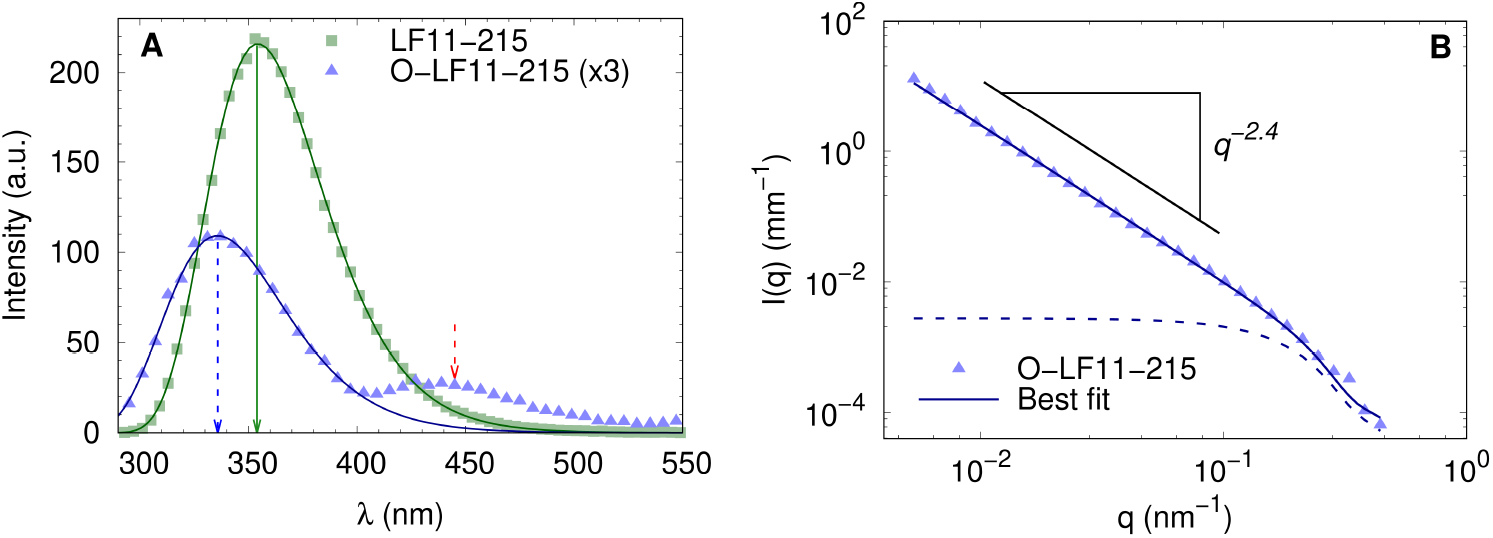
(**A**) Trp fluorescence data of LF11-215 (green sqares) and O-LF11-215 (blue triangles) at 100 *μ*g/ml (LF11-324 are not shown to avoid redundancy). Data were fitted with **Equation 4**. Arrows mark the maxima positions of the fluorescence and phosphorescence bands. (**B**) SAXS data of O-LF11-215 at 400 *μ*g/ml. The fit was performed with **Equation 5**.

USAXS/SAXS data in the low *q*-range (*q_min_* ~0.005 nm^−1^) exhibited a featureless decay of intensity with a slope of *f* = 2.4. This slope value is typical for mass fractals, i.e. highly branched objects with high surface-to-volume ratio, while *q_min_* suggests a minimum aggregate size of ~ 2*π*/*q_min_* >1 *μ*. Furthermore, a Guinier term is needed to fit the shoulder at about *q* = 0.2 nm^−1^ corresponding to an average radius of gyration *R_g_* ≃ 10 nm. Note that this feature also does not vanish for different choices of scaling constants for background subtraction. Interestingly, this value is way too high to describe O-LF11-215 monomers, whose expected radius of gyration would be < 1 nm. Possibly, peptide monomers create smaller aggregates of mean size *R_g_* ≃ 10 nm, which in turn form an heterogeneous and branched supramolecular structure with the characteristics of a mass fractal.

## Appendix 2 Estimating the scattering contribution of OMVs

The prefactor of the scattering contribution from extracellular, independent objects used in **Equation 6** is 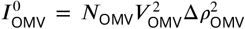, where *N*_OMV_ is the number of OMVs, *V*_OMV_ is the volume of an OMV and Δ*ρ*_OMV_ is average the SLD difference to the buffer. This definition of forward scattering (per single cell) is valid for every non-interacting object. Hence, to validate the assumption that this scattering contribution is related to OMVs, it is interesting to calculate possible *N*_OMV_, *V*_OMV_ and Δ*ρ*_OMV_ values. Note that even if modelling OMVs as homogeneous spheres appears as a crude first order approximation, *R*_OMV_ can be associated to its radius (within ~10% confidence). Assuming the same lipid asymmetry as in the outer membrane, the inner leaflet of OMVs can be mimicked by a 3:1 mole mixture of palmitoyl-oleoyl-phosphatidylethanolamine (POPE) and palmitoyl-oleoyl-phosphatidylglygerol (POPG), respectively (***De Siervo, 1969***; ***Lohner et al., 2008***; ***Leber et al., 2018***). The SLD membrane profiles of these lipids have been thoroughly investigated (***Kučerka et al., 2012, 2015***). The outer leaflet might instead be dominated by LPS, whose lipid A possesses about 6 short C14:0 chains (***Kim et al., 2016***), and the polar region can be approximated as two PG units, in terms of molecular volume and SLD. In addition, LPS inner and outer core volumes and SLDs can be calculated from ***Heinrichs et al.*** (***1998***) and ***Müller-Loennies et al.*** (***2003***), neglecting O-antigen chains for simplicity. Gathering all this information, similarly to the membrane structure estimation in ***Semeraro et al.*** (***2021***), the vesicles would have a membrane thickness of 4.1 nm and an average SLD of 1.1×10^−3^ nm^−2^ (volume-weighted averages). The lumen of OMVs can be quite diversely composed (***Beveridge, 1999***). We tentatively assigned the SLD of the periplasmic space of the end-state system, i.e. 9.68×10^−4^ nm^−2^.

Together with a buffer SLD of 9.47×10^−4^ nm^−2^, a measured 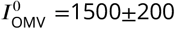 nm and a *R*_OMV_ =15.4±0.6 nm, for instance, then leads to the estimate *N*_OMV_=1200±400 and a total lipid surface (inner and outer leaflets of all OMVs) of (4±2)×10^6^ nm^2^.

## Appendix 3 Tryptophan fluorescence

**Appendix 3 Figure 1.**
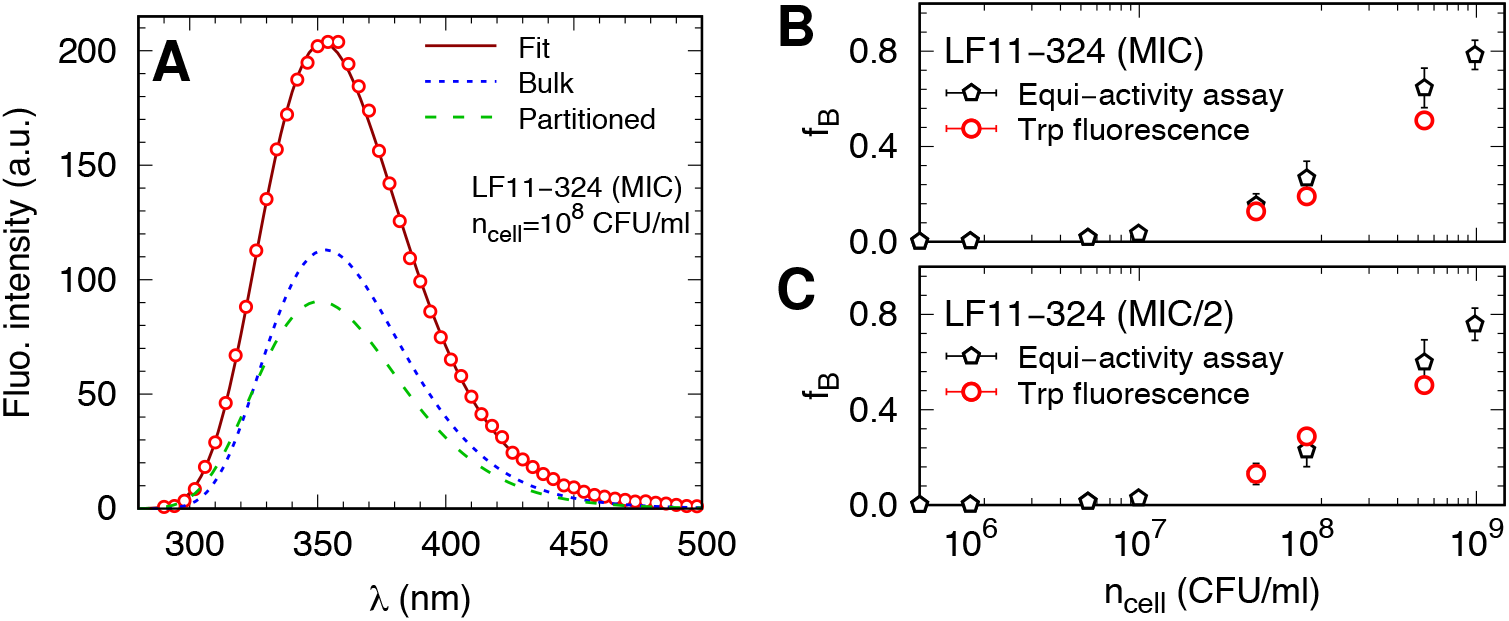
(**A**) Example of Trp fluorescence analysis in LF11-324 systems. The solid line is the best fit and the dotted and dashed lines represent the Trp emissions from peptide in bulk and cell-associated, respectively. Data were fitted with **Equation 4**. (**B-C**) Comparison between *f_B_* values obtained from the Trp fluorescence analysis (red dots) and the equi-activity analysis from the susceptibility assay (black diamonds).

The native fluorescence of the single Trp residue present in LF11 peptides was exploited to validate the partitioning investigation through the equi-activity analysis. For every system, emission signals from partitioned peptides exhibited a weak blue-shift, with 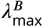 values in the range 346–354 nm for LF11-215, and 340–350 nm for LF11-324. Γ_*B*_ values showed no significant variations, instead. Interestingly, these values are consistent with a scenario in which a significant amount of partitioned peptides are heterogeneously dispersed in a polar environment and in a configuration allowing full dynamics of the Trp residues (***Burstein and Emelyanenko, 1996***; ***Ladokhin et al., 2000***). This is consistent with the 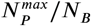 values estimated via *ζ*-potential measurements, suggesting that a relevant portion of partitioned AMPs are still in monomeric state in the cytosol

Given that *f_B_* = *n*_cell_*N_B_*/(*N_A_*[*P*]), *f_B_* values extracted via Trp fluorescence were compared with those obtained from the antimicrobial activity assays at [*P*] = MIC and MIC×0.5 (LF11-215 data not shown). These two independent methods gave comparable *f_B_*, confirming the validity of *N_B_* values.

## Appendix 4 Statistics of bacterial inhibition

Assuming that the AMP-induced delayed bacterial growth is entirely due to a lower number density of survived cells (***Marx et al., 2021b***), the inhibited fraction of cells, *ϕ*_IG_, as a function of peptide and cell concentrations was fitted with a heuristic approach. Specifically, we used the sigmoidal Gompertz function

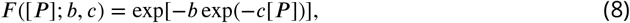

where [*P*] is the total peptide molar concentration, and *b* and *c* are related, respectively, to the position and width of the sigmoidal.

*F*([*P*]; *b*, *c*) can be associated to a cumulative distribution function (CDF), that is, it measures the probability of finding a certain number (or fraction) of inhibited cells at a given peptide concentration. This allows to derive the probability density function (PDF) by calculating the derivative, *f*([*P*]; *b*, *c*) = ∂*F*([*P*]; *b*, *c*)/∂[*P*] = ∂*ϕ*_IG_/∂[*P*], as well as the inverse CDF, *F*^−1^(*ϕ*_IG_; *b*, *c*), which maps *ϕ*_IG_ values to the inhibitory peptide concentrations IC_*x*_, where *x* is the corresponding inhibited bacterial percentage; by definition, IC_99.9_ ≡ MIC.

In addition, the set of *b* and *c* values as a function of *n*_cell_ can be interpolated to obtain, for example, a continuous trend of IC_*x*_ as a function of *ϕ*_IG_.

**Appendix 4 Figure 1.**
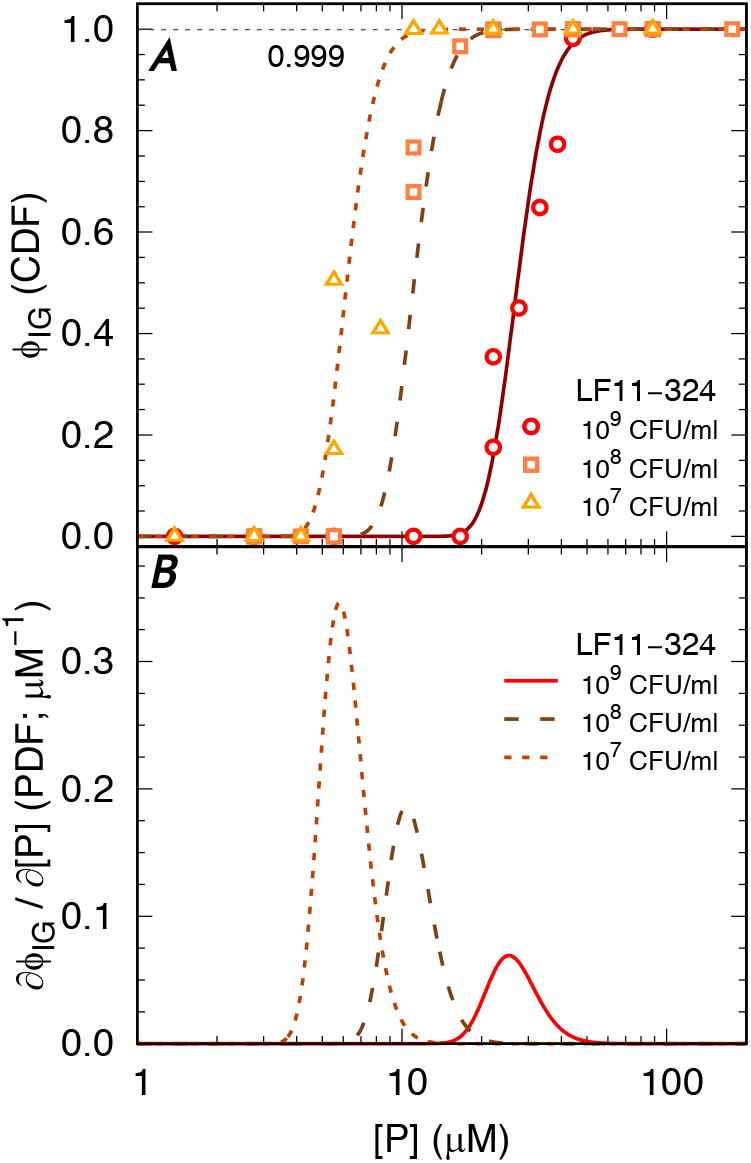
(**A**) Selected *ϕ*_IG_ data for LF11-324 and corresponding fits with the Gompertz function. (**B**) Corresponding PDFs.

---

**Figure 1–Figure supplement 1.**
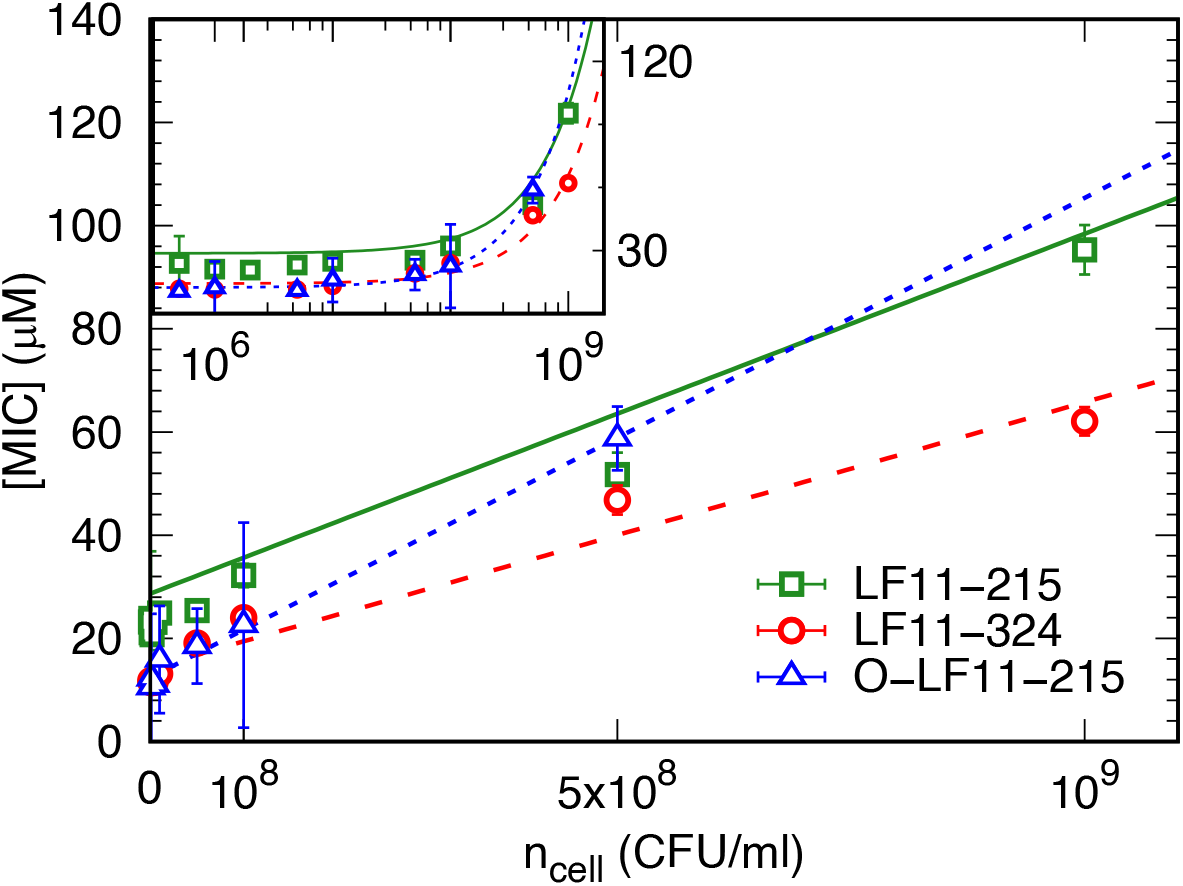
MIC values as a function of *n*_cell_ for LF11-215 (green squares), LF11-324 (red dots) and O-LF11-215 (blue triangles) and best fits using **Equation 1** and **Equation 2**.

**Figure 1–Figure supplement 2.**
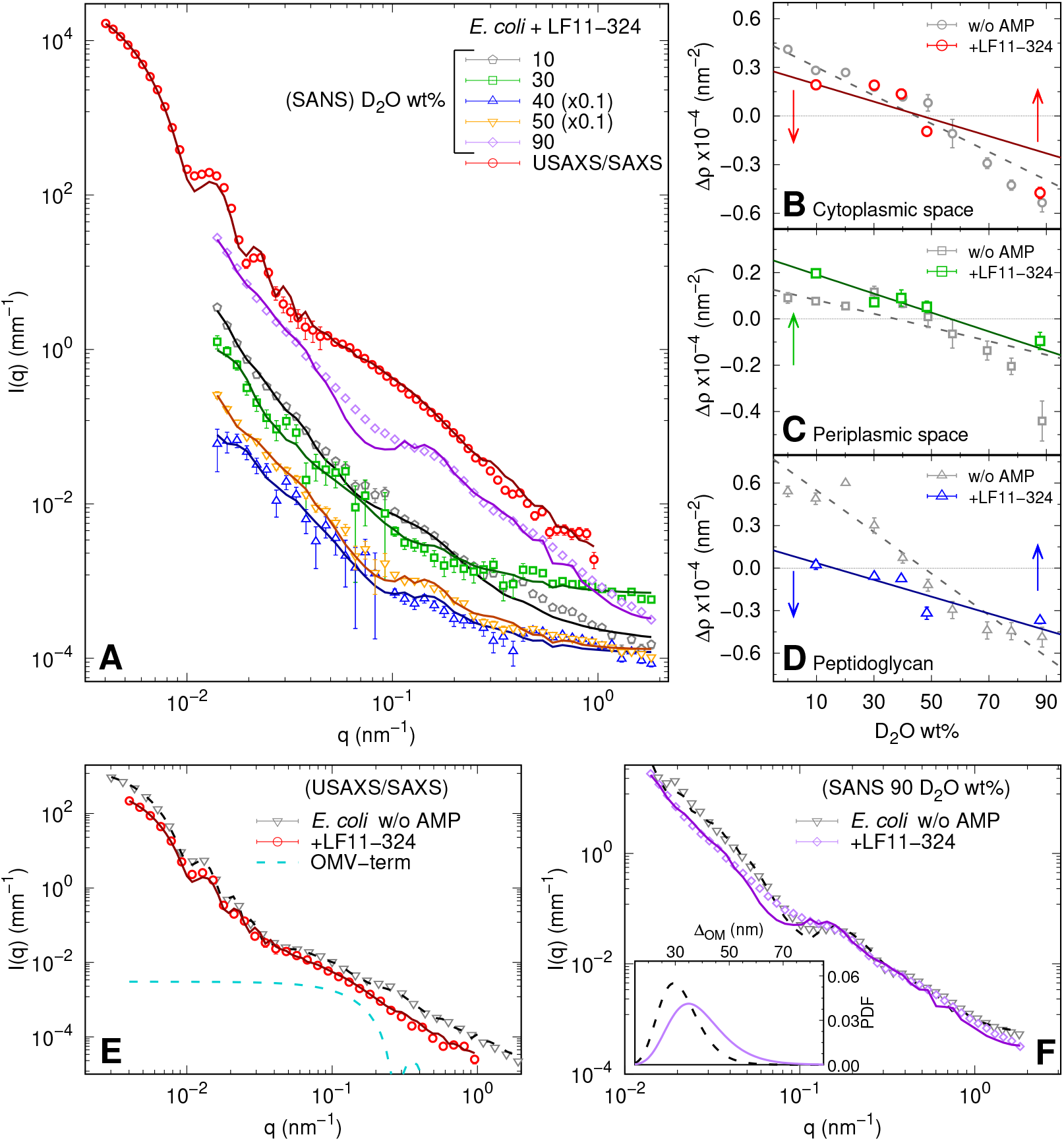
(**A**) X-ray and neutron scattering data of bacterial systems after one hour incubation with LF11-324 (end-states) at the MIC (SANS) and 1.2×MIC (SAXS). Lines are the best fits using **Equation 6**. (**B-D**) Scattering length contrasts as a function of D_2_O concentration (wt%) in the medium for the cytoplasm, Δ*ρ*_CP_, periplasm, Δ*ρ*_PP_, and peptidoglycan layer, Δ*ρ*_PG_. Gray symbols are the values reported in absence of peptides [data adapted from ***Semeraro et al.*** (***2021***)]. Solid and dashed lines correspond to linear regressions. (**E**) Comparison between SAXS curves from bacterial end-states and a reference sample without peptides [data adapted from ***Semeraro et al.*** (***2021***)]. The blue dashed line represents the additional *I*_OMV_ term whereas the fits refer to **Equation 6**. (**F**) Comparison between SANS curves from bacterial end-states and a reference sample without peptides [data adapted from ***Semeraro et al.*** (***2021***)] in the case of 90 D_2_O wt%. Lines are the best fits with **Equation 6**. The inset shows the log-normal probability distribution function (PDF) of the inter-membrane distance with (dashed purple line) and without LF11-324 (solid black line). (**E-F**) The intensities are on absolute scale and normalized by the cell concentration.

**Figure 1–Figure supplement 3.**
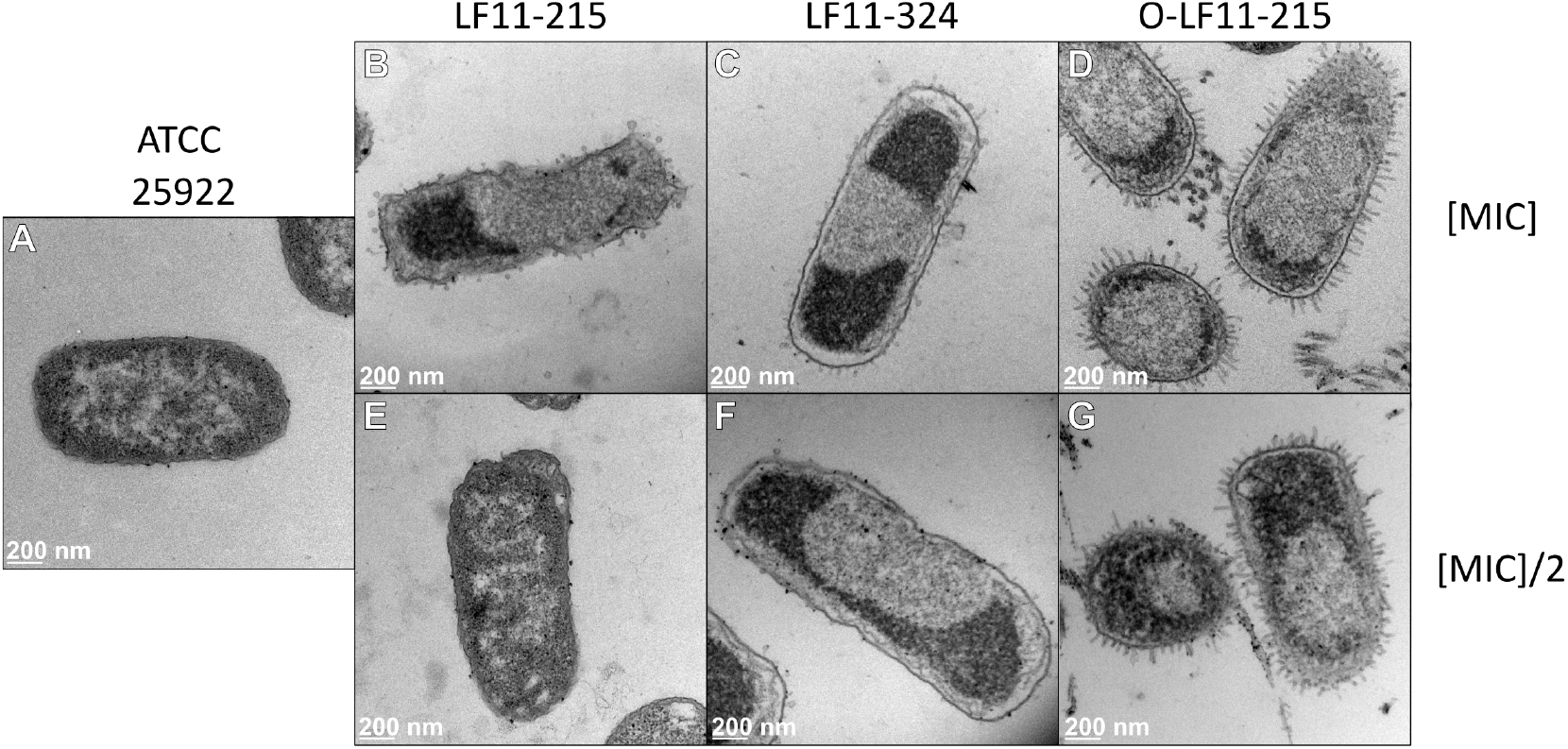
TEM images of *E. coli* ATCC (**A**) and end-states in the presence of LF11-215 (**B,E**), LF11-324 (**C,F**) and O-LF11-215 (**D,G**). All systems were probed at the MICs (**B,C,D**) and half the MICs (**E,F,G**).

**Figure 2–Figure supplement 1.**
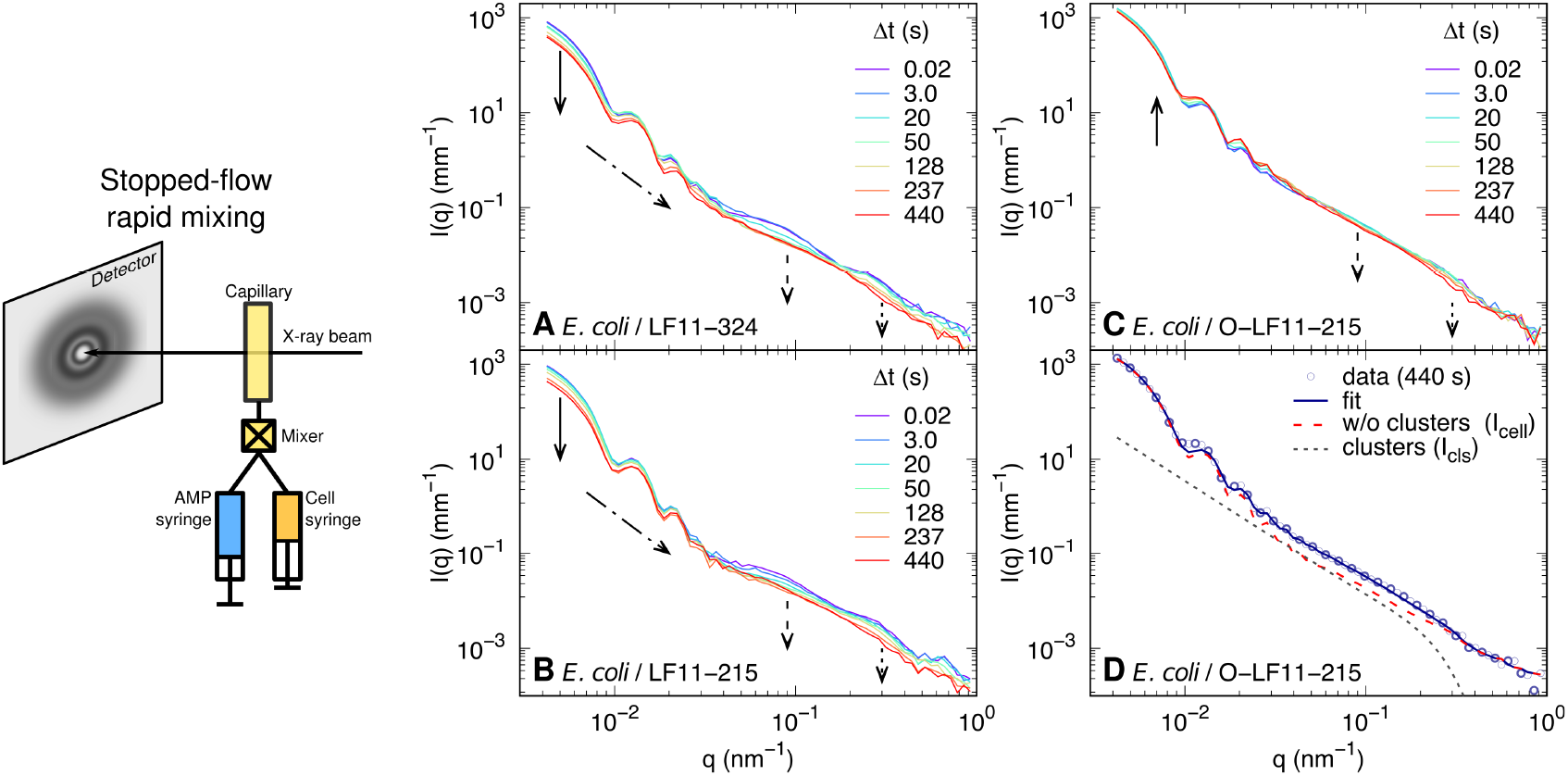
Schematic of the stopped-flow rapid mixing set-up used for USAXS/SAXS experiments and kinetic changes of USAXS/SAXS curves of bacterial samples upon mixing with LF11-324 at MIC×1.2 (**A**), LF11-215 at MIC×1.6 (**B**) and O-LF11-215 at MIC×1.7 (**C**). The arrows highlight the most significant variations of intensity, such as the decrease of forward scattering (**A-B**);the evolution of the low-<u>q</u> oscillation profile (**A-C**);the disappearance of the feature at *q* ~ 0.3 nm^−1^ (**A-C**);the decrease of *I*_cell_ (**A-B**) and of *I*_clu_ (**C**) at *q* ~ 0.1 nm^−1^. (**D**) Example of the combination of *I*_cell_ and *I*_clu_ in the case of applying O-LF11-215 (see **Equation 6**).

**Figure 2–Figure supplement 2.**
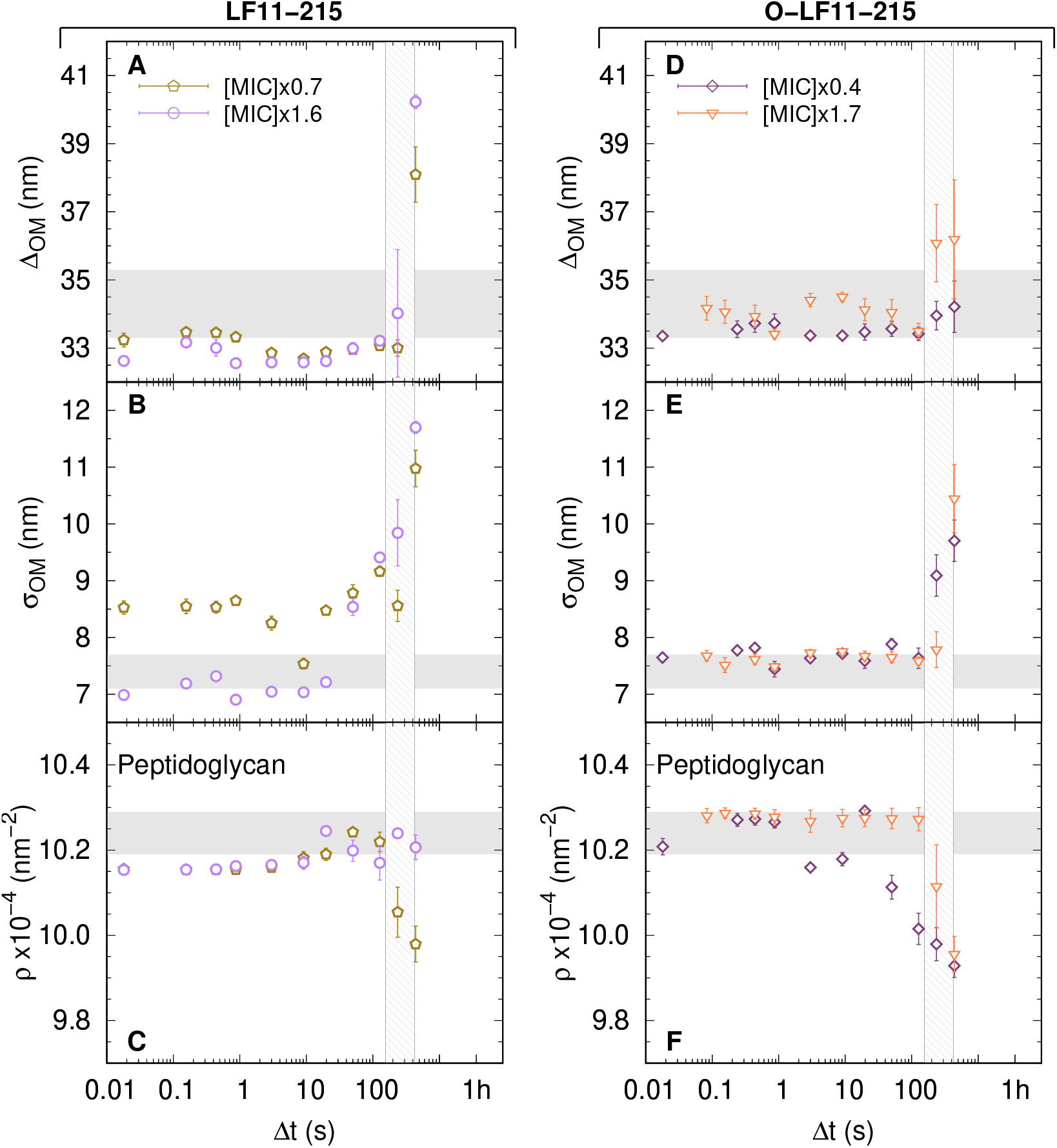
(**A-C**) Kinetics of the adjustable parameters upon mixing with two concentrations of LF11-215. These are, the intermembrane distance (~ periplasm thickness) (**A**), its deviation from the average value (**B**), and the peptidoglycan SLD (**C**). (**D-F**) Kinetics of the adjustable parameters upon mixing with two concentrations of O-LF11-215. The parameters are the intermembrane distance (~ periplasm thickness) (**D**), its deviation (**E**), and peptidoglycan SLD (**F**). Thick gray bands mark the degree of confidence from bacterial systems w/o peptides [see **Table 1** and ***Semeraro et al.*** (***2021***)]. The vertical gray grid in (**A-F**) is an approximated, concentration-independent time range during which the cell-wall damage occurs.

**Figure 2–Figure supplement 3.**
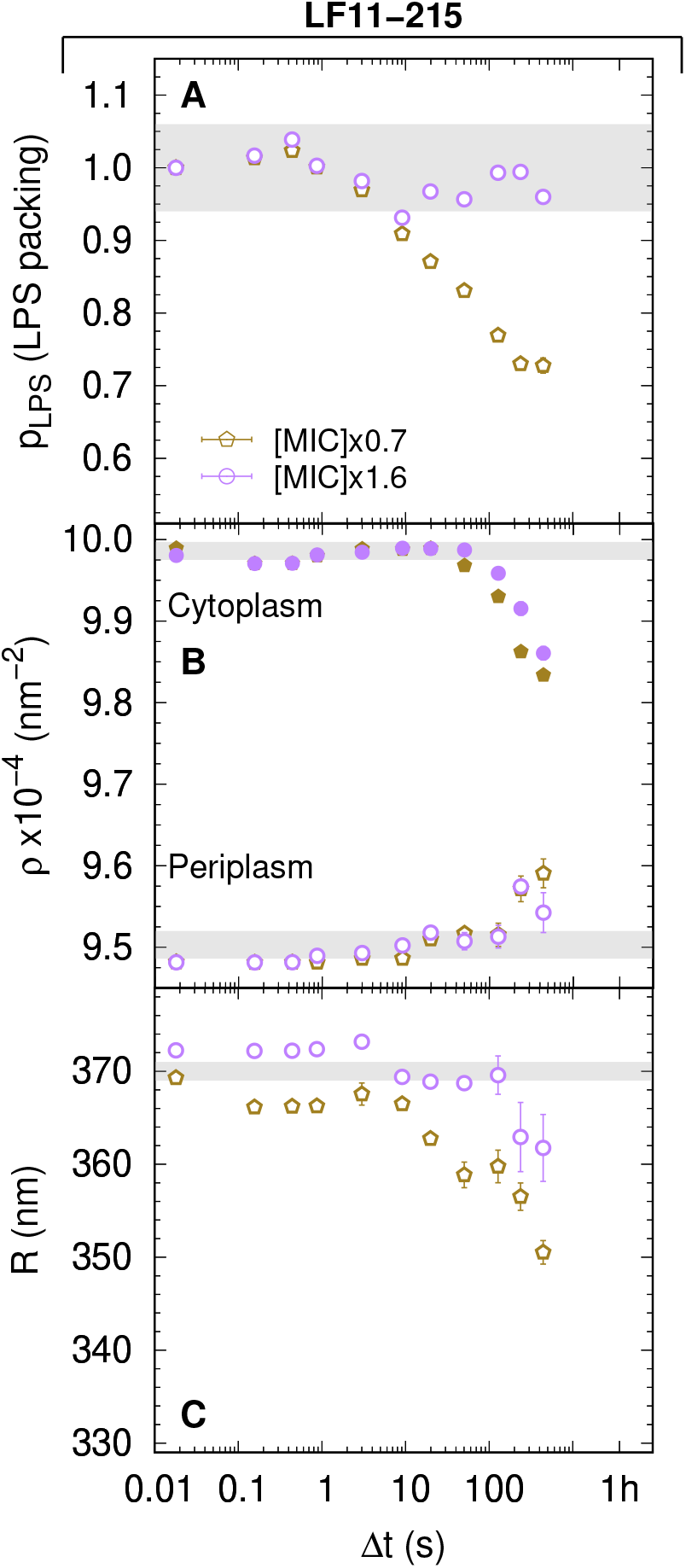
(**A-C**) Kinetics of the adjustable parameters upon mixing with two concentrations of LF11-215. These parameters are the LPS packing (**A**); the cytoplasm and periplasm SLDs (**B**); and the minor radius of the cell (**C**). Thick gray bands mark the degree of confidence from bacterial systems w/o peptides [see **Table 1** and ***Semeraro et al.*** (***2021***)], except for (**C**), where they refer to the average of the current cell radii at Δ*t* = 0.0175 s.

**Figure 4–Figure supplement 1.**
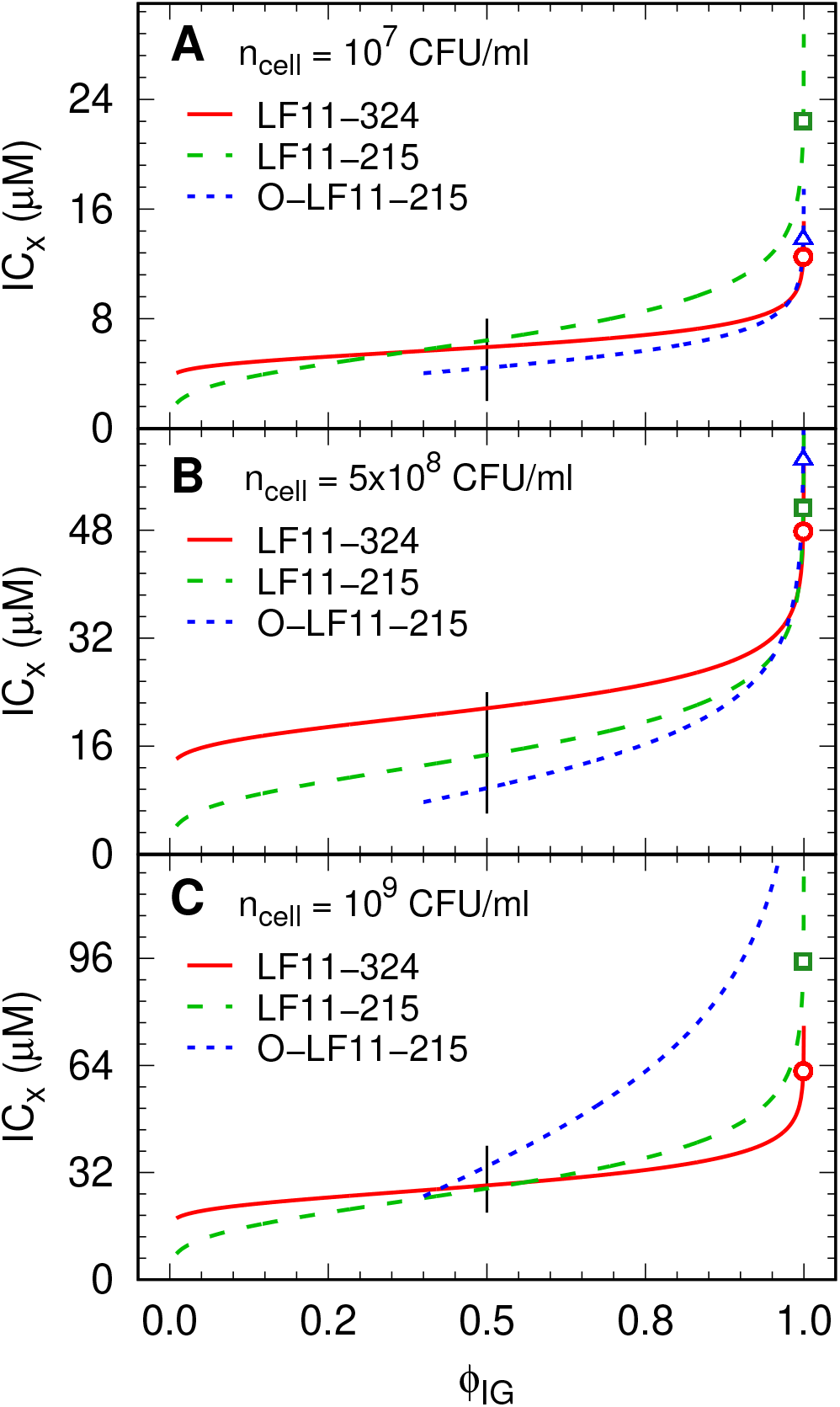
(**A-C**) Inhibitory concentration (IC_*x*_) as a function of inhibited fractions *ϕ*_IG_ [inverse CDF, *F*^−1^(*x*; *b*, *c*)] for different peptide and cell concentrations. Low *ϕ*_IG_ values for O-LF11-215 were not accessible due the high noise-to-signal ratio. Symbols mark the MICs for LF11-324 (circles), LF11-215 (squares) and O-LF11-215 (triangles), and black lines mark the range of IC_50_. The level of confidence is not displayed for sake of clarity. IC_*x*_ values have an about 10% relative error.

**Figure 5–Figure supplement 1.**
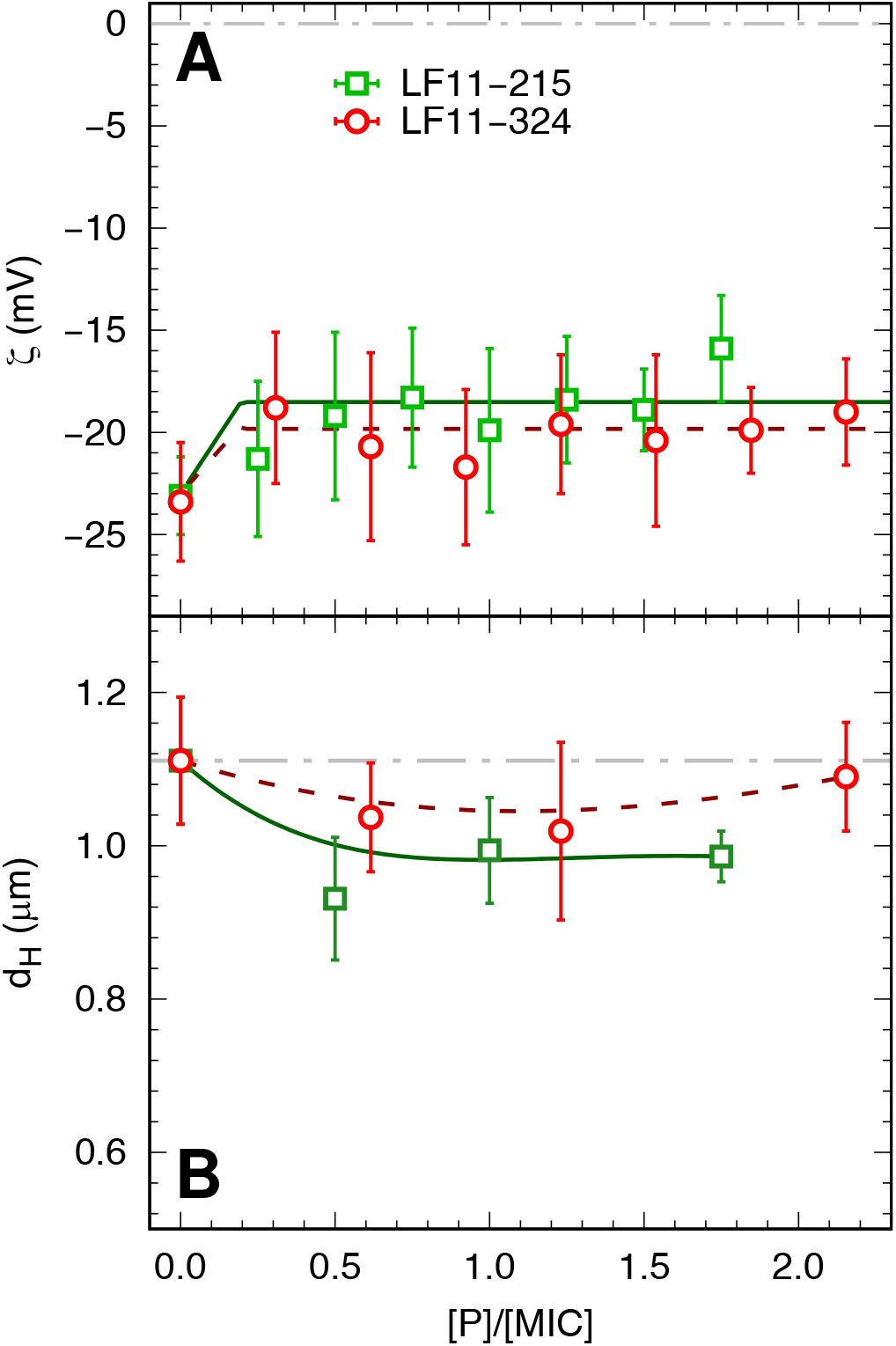
*ζ*-potential (**A**) and size (**B**) measurements as a function of peptide concentration (normalized by the respective MICs of LF11-215 and LF11-324 peptides). Lines are guides for the eye.

**Figure 5–Figure supplement 2.**
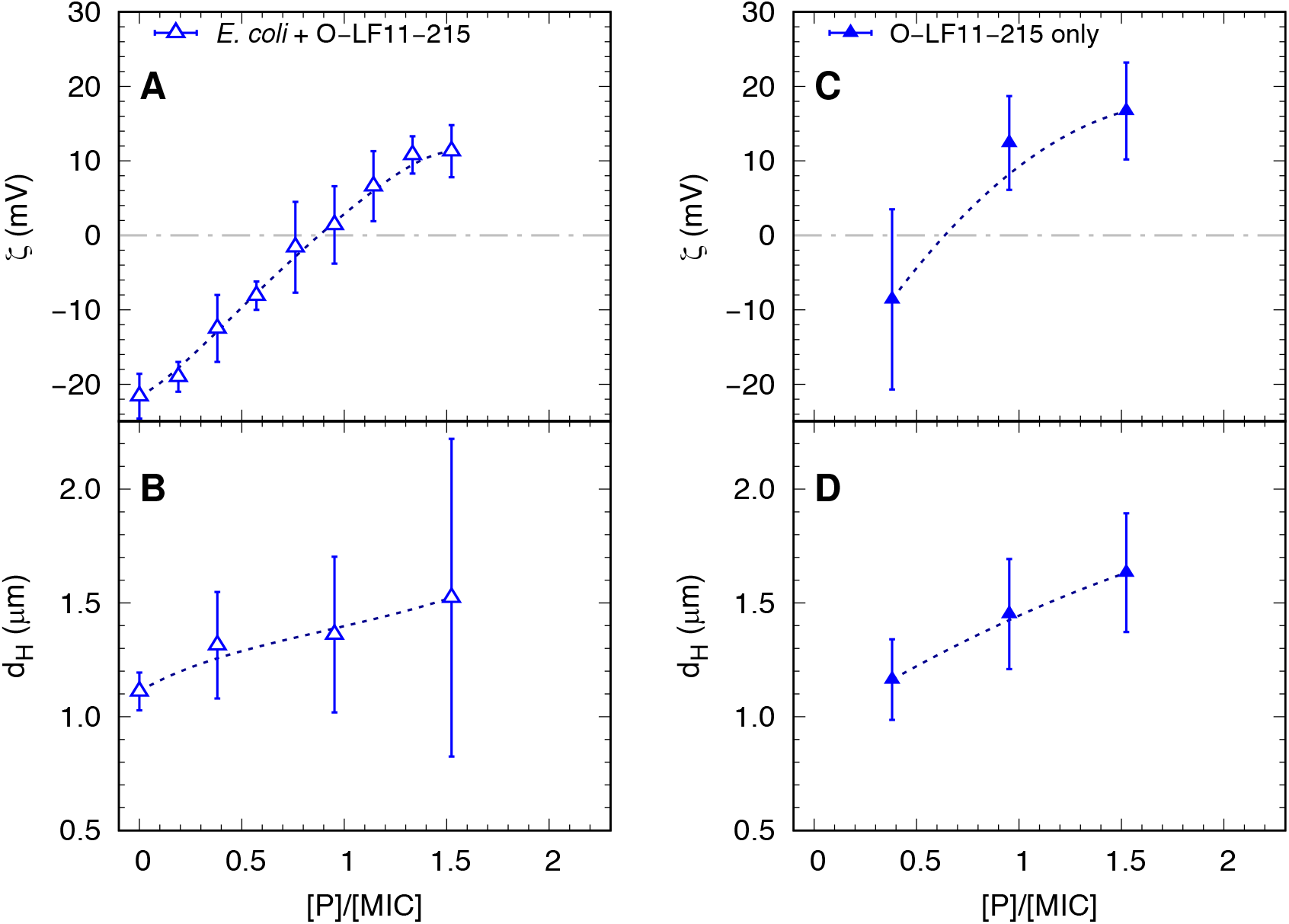
Comparison between *ζ*-potential (**A,C**) and size (**B,D**) measurements of O-LF11-215 AMP alone and mixed with *E. coli* as a function of peptide concentration (normalized by the MIC of O-LF11-215).

